# Vital Insights into Prokaryotic Genome Compaction by Nucleoid-Associated Protein (NAP) and Illustration of DNA Flexure Angles at Single Molecule Resolution

**DOI:** 10.1101/2020.09.11.293639

**Authors:** Debayan Purkait, Debolina Bandyopadhyay, Padmaja P. Mishra

**Author notes:** Both the authors have contributed equally (Debayan Purkait, Debolina Bandyopadhyay).

## Abstract

Integration Host Factor (IHF) is a heterodimeric site-specific nucleoid-associated protein (NAP) well known for its DNA bending ability. The binding is mediated through the narrow minor grooves of the consensus sequence, involving van der-Waals interaction and hydrogen bonding. Although the DNA bend state of IHF has been captured by both X-ray Crystallography and Atomic Force Microscopy (AFM), the range of flexibility and degree of heterogeneity in terms of quantitative analysis of the nucleoprotein complex has largely remained unexplored. Here we have monitored and compared the trajectories of the conformational dynamics of a dsDNA upon binding of wild-type (wt) and single-chain (sc) IHF at millisecond resolution through single-molecule FRET (smFRET). Our findings reveal that the nucleoprotein complex exists in a ‘Slacked-Dynamic’ state throughout the observation window where many of them have switched between multiple ‘Wobbling States’ in the course of attainment of packaged form. A range of DNA ‘Flexure Angles’ has been calculated that give us vital insights regarding the nucleoid organization and transcriptional regulation in prokaryotes. This study opens up an opportunity to improve the understanding of the functions of other nucleoid-associated proteins (NAPs) by complementing the previous detailed atomic-level structural analysis, which eventually will allow accessibility towards a better hypothesis.

## Introduction

The folding of DNA inside the cell is a prerequisite for survival in all domains of life. Genetic analysis of the genes assigned in PEDANT^1^ reveals that 2-3% of the prokaryotic genome and 6-7% of the eukaryotic genome encodes for various DNA binding proteins that are associated with genome compaction.^2^ In eukaryotes, this process is governed by histone and other chromatin remodelling factors in a highly organized and sequential manner. Unlike eukaryotes, where the genome is organized by repeated arrays of structural units, the bacterial genome adopts a range of different conformations by the nucleoid-associated proteins (NAPs).^3^ NAPs help in genome compaction by either wrapping or bending the DNA as oligomers in both specific (sequence-specific or structure-specific) and non-specific manner.^4^ The specific binding of NAPs is associated with gene regulation whereas the non-specific binding has been attributed primarily to genome compaction and stress response.^5^ Though the histones and other chromatin remodellers have been widely studied both *in vivo* and *in vitro*, less focus is been given on the NAPs till date, with a few exceptions.

Integration Host Factor (IHF) is one of the heterodimeric NAPs, having two 10 kDa subunits, α and β with approximately 30 percent sequence homology, is well known for its DNA binding and bending ability.^6;7^ It was identified as a host factor for lysogeny by bacteriophage λ, where it directly participates in the recombination reaction by binding with the region of viral DNA that takes part in cross-over. One of the tightest binding sites of IHF is the H’ site (5’TATCAA3’), a subset of the 250 base pairs long attP region of phage λ.^8; 9^ It falls into the category of minor-groove intercalating protein and known to reverse the direction of the helical axis providing a pseudo-two fold symmetry to the DNA by inducing two kinks within a very short distance (9 base pairs apart).^6^ The action is very similar to the U-shape bending induced by the nonspecific architectural histone-like protein, HU, as seen by the NMR and X-ray Crystallography studies.^10^ Though IHF and HU possess closely similar structures and share several regions of conserved homologies, unlike HU, IHF binds to DNA both specifically during site-specific recombination, DNA replication and transcription and non-specifically, in its role as a DNA compaction protein.^11; 12^

The two arms of β-ribbons of IHF curl like a clamp to both sides of the DNA in the vicinity of H’ region stabilizing the kink in the DNA, where one proline from the tip of each β ribbon intercalates between the base pairs of the DNA. The specific complex between the minor groove of DNA with IHF involves van der-Waals interaction with very similar H-bonding patterns offered by different bases, and electrostatic contacts with the sugar-phosphate backbone.^6^ Prior to binding with the H’ site, IHF performs a quick search via rapidly diffusive ‘indirect readout’ mechanism and spots the narrow minor groove formed by an A-tract region (5’AAAAAA3’) present close to the H’ sequence before switching to a slow diffusive ‘direct readout’ mechanism to bind to the DNA through a conformational change.^13; 14; 15^ Although previous studies involving nanosecond temperature-jump measurements coupled with Stopped-flow and FRET have indicated the existence of short-lived transient intermediate states in the H’-IHF complex, the variations among them in terms of both population distribution and degree of bending are not quantified. ^16; 17^ The binding of protein engenders considerable architectural changes in the DNA leading to local deformation, twisting or bending of the B-DNA structure, and makes it feasible to bring several regulatory elements closer and carry out gene regulatory processes.^18; 19^ The genome reorganizing property of IHF is reported to play a crucial role in numerous cellular activities like replication,^3; 20; 21; 22^ transcription and gene regulation,^23; 24^ bacterial growth cycle,^25; 26^ distribution of transposable elements,^27; 28^ folding of DNA into viral capsids,^29^ and genome compaction.^23^ Recently, IHF has gained significant attention from both the medical and pharmaceutical community due to its active role in the formation and stabilization of biofilms in cystic fibrosis.^30; 31^The reports so far provide significant insights into the finely tuned interactions between IHF and its cognate sites that persuade bending in DNA by reconciling quantitative information on the dynamic equilibrium between different DNA conformations (kinked or not kinked) and correlates the receptivity on DNA sequence and deformability.

Though the nature of the distortions in DNA as a consequence of complexation with IHF has revealed at atomic resolution, the temperament of the conformational plasticity of DNA in solution needs further investigation. The works so far have not been able to evaluate the structural alternations, in particular, the degree of compaction on DNA induced by IHF. This strengthens the fact that classical techniques used in molecular biology are designed for monitoring relatively strong DNA-binding sites and are not effective in appraising the contribution of nonspecific or low-affinity interactions and to overcome this, various other sophisticated experimental techniques like chromatin immunoprecipitation (ChIP), in situ Hi-C, pulsed-EPR double electron-electron resonance (DEER), and three-color FRET with alternating-laser excitation (ALEX) are progressively being used to probe the dynamic nature of biological systems.^32; 33; 34^ In this study, we have employed a single-molecule Förster Resonance Energy Transfer (smFRET) micro-spectroscopic method to compassionate the dynamic fluctuations of the nucleic acid in IHF bound H’ DNA complex (Figure 1). One of our prime concerns is to investigate the elasticity of the single dsDNA bound to IHF molecule to see if the nucleoprotein complex exhibits inflexible kinks or flexible hinges, by avoiding the ensemble averaging of quantized information. We have also attempted to elucidate the dynamical properties acquired by the local A-tract region upon inducement of curvature to the dsDNA due to IHF binding considering the sensitivity in monitoring the change in FRET value during conformational transposition in DNA, we have labelled different regions of the DNA to figure out the region of the nucleic acid that demonstrates effective dynamicity upon protein binding. The DNA sequence used in this study has an interruption of asymmetric A-tract units that are reported to be dynamically stiffer than G/C rich sequences. In response to local and global conformational changes and induced bending of DNA, the A-tract region gains flexibility mostly in favour of promoting compaction of the genome. The intrinsic change in structural features due to the presence of A-tract sequence causes various gene regulatory processes to get kick-started that includes recruitment of transcription and replication factors and promotion of gene expression.^35^

**Figure 1.**
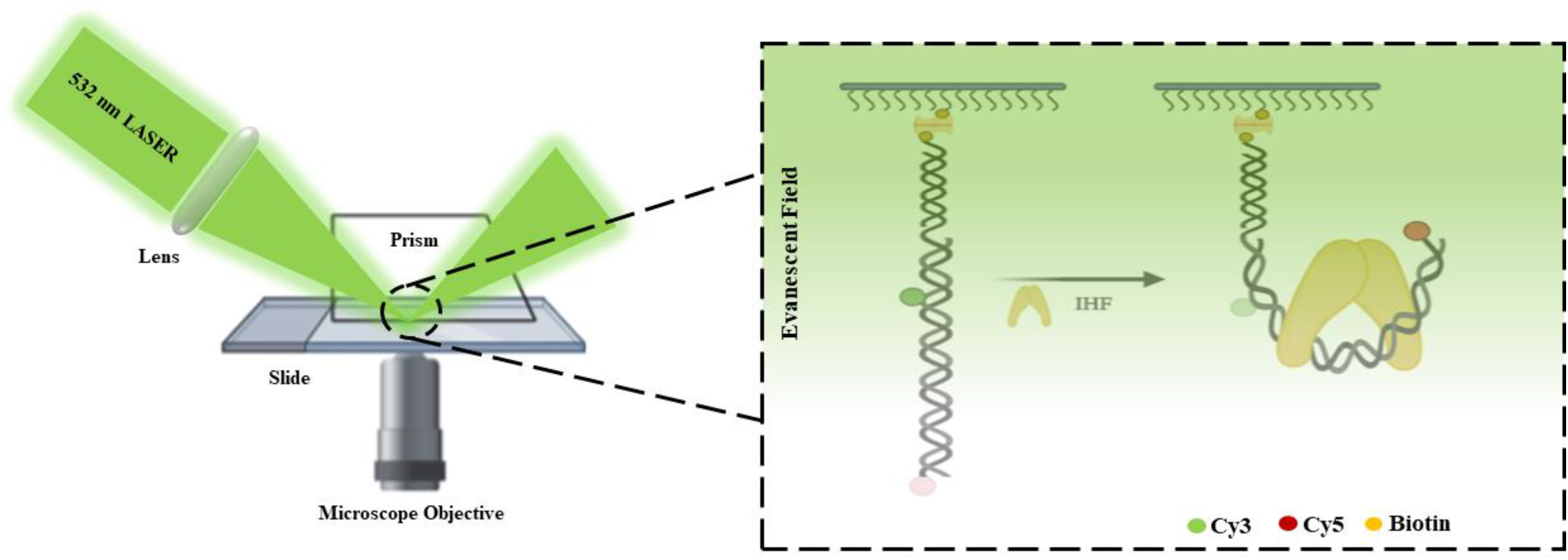
Schematic representation of Prism-type Total Internal Reflection Microscopy (TIRF) based smFRET. The figure was created with BioRender.com.

Additionally, it has been observed that linking the individual α and β subunits of the wild-type IHF (wtIHF) by a short peptide linker enhances the stability of the protein (single-chain IHF with 204 amino acids), as biochemically the wild-type IHF (wtIHF having 193 amino acids) is not very stable *in vitro* and prone to rapid aggregation.^36^ Previous biochemical studies have reported that there are no significant changes in the activity of the protein due to this modification. This provoked us to think why nature would favour a less stable variant and since it did, it needs to be validated if there are reasons. Thus, we have compared the bending dynamics of single-chain IHF (scIHF) and wild-type IHF (wtIHF) from *E*.*coli*at single molecular resolution by allowing them to bind with the same pair of labelled dsDNA. Finally, from the smFRET measurements, we have come up with a conceptual model to quantify and illustrate the different sequential bending of the DNA curvature through a range of ‘Flexure Angles’^37^ by the two variants of IHF.

## Results

### Characterization of conformational plasticity of dsDNA terminals due to IHF binding

To elucidate the dynamics of DNA bending, a 35 base pair long DNA is used in this study carrying 6 adenine repeats and H’ binding site for IHF and is site-specifically labelled with Cy3 and Cy5 FRET pairs at the terminal nucleotides named as Construct 1 (Supplementary Figure S1a and Table S1). IHF binding causes architectural changes in the DNA, leading it to bend and hence the ends approach each other (Figure 2a). The depreciated end to end distance is quantitatively monitored by the FRET efficiency values upon applying suitable bleeding correction and background subtraction, as described elsewhere.^38^ In the absence of protein, the DNA only illustrated a single low FRET state (0.1) without any noticeable change in FRET efficiency until photo-bleaching. It is worth mentioning here that the sequence of the oligonucleotides used for the experimental purpose are smaller than their corresponding persistent length, keeping in mind that they behave as an isotropic rod. This is also supported by the FRET values obtained with population maxima of the histogram at 0.1 (Supplementary Figure S2a, Figure 2f). Bending in the DNA would certainly increase the FRET Efficiency and hence, the population of the histogram shows a subtle shift towards higher FRET state upon interaction with either wtIHF or scIHF. However, this resultant FRET value is not as high as it should have been with the formation of the nucleoprotein complex captured in the crystal structures (Supplementary Figure S3a and Table S1). This ambiguity in the observed value indicates that upon interaction with the protein the arms of the DNA frays to wrap itself over the protein in the course of attaining a packaged form instead of attaining a quick sharply bent conformation. For analysis purposes, the molecules are separated based on if they possess a single FRET state till photo-bleaching or exhibit multiple conformational states and separate histograms are generated. Around 60% of DNA molecules present changed their dynamical states by transitioning between multiple conformers upon binding with scIHF, whereas the same for wtIHF is ∼24% (Figure 2c). Complexation with wtIHF leads multiple state transitions with two distinct FRET values, one at 0.1 that corresponds to the open DNA conformation and the other with a slightly close DNA ends having *e*_*fret*_ around 0.25 (Figure 2f). The time traces of the molecules showing dynamic transition fluctuation in their behaviour were subjected to two sate Hidden Markov Model (HMM) Analysis and the transition matrix so found as output is extrapolated to get the temporal kinetic rates of transition. The rate of transition from the unbent state to the partially bent state is found to be faster (*k*_*12*_=2.6 s^-1^) compared to the reverse process (*k*_*21*_=1.2 s^-1^, Figure 4a). However, the mean dwell time, calculated manually as the time spent by molecules at these two distinct states, turned out to be almost comparable (*τ*_*1*_ = 10.9 seconds for *e*_*fret*_ = 0.1 and *τ*_*2*_ = 11.7 seconds for *e*_*fret*_ =0.25 FRET states, Figure 4b).The peak positions in the Gaussian fitted histogram for the scIHF-DNA complex also shows very close values of FRET Efficiency (0.12 and 0.23), however, they exemplify a difference in population distribution at the two dynamically equilibrium states (Figure 2f). Moreover, the efficiency histogram is narrower, indicating that the population possesses the property of less dynamic averaging as compared to the DNA-wtIHF complex. The broadening of the histogram for the wtIHF-DNA portrays higher-order molecular heterogeneity, less integrity, and more complexities in the same micro-environment. The rates of transition also followed a very similar trend as observed for wtIHF, with a marginally higher dwell time of 36.8 seconds at the unbent state and 26.7 seconds for the other state (Figure 4c, 4d). The static conformation possessing molecules shows a peak at 0.22 FRET Efficiency and a very small population at 0.5 when subjected to wtIHF binding (Figure 2f). However, there is a difference in the conformational states of the DNA when bound to scIHF with two FRET states of 0.19 and 0.30, both illustrating bent conformation (Figure 2f).The multiple dynamic states as observed in the FRET measurements fit closely with physiological circumstances keeping view of the fact that even a short and rigid DNA sequence develops the property of acquiring flexibility while acting as a regulatory unit or when needs to be packaged.^39^

**Figure 2.**
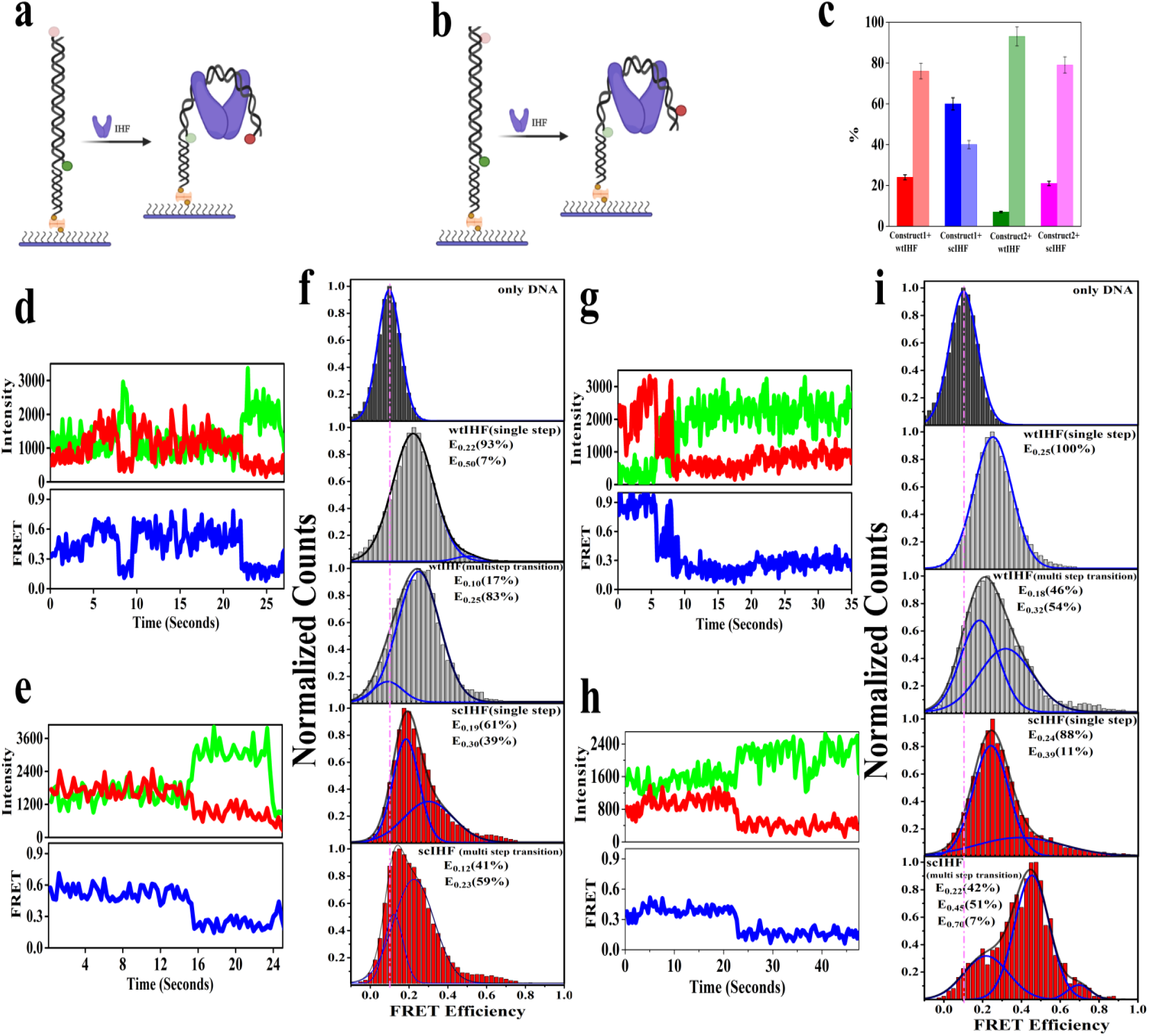
IHF induced bending of Construct 1 and 2. **(a)** and **(b)** corresponds to the representation of the cartoon model of experimental set-up with Construct 1 and 2 of DNA respectively. **(c)** Bar plot characterizing relative fraction of molecules exhibiting dynamic FRET states (dark-colored bars) and molecules existing in a constant FRET state throughout until photo-bleaching. **(d)**and**(e)**is the representative time traces and FRET efficiency plots of Construct 1 of DNA when exposed to the proteins wtIHF and scIHF respectively. **(f)** FRET Histograms illustrating efficiency traces of DNA Construct1 (top), DNA when exposed to wtIHF, and exhibiting single FRET state followed by multiple state representing molecules. The last two boxes of FRET efficiency histogram show DNA when getting bound to scIHF with static and dynamic molecules being segregated. **(g)** and**(h)**are FRET traces and corresponding efficiency plots of DNA Construct 2 when subjected to bind with wtIHF and scIHF respectively. **(i)** FRET Histograms of Construct 2 naked DNA, also with scIHF and wtIHF proteins molecules being segregated in terms of single state exhibiting and multiple dynamic state exhibiting behaviors.

**Figure 3.**
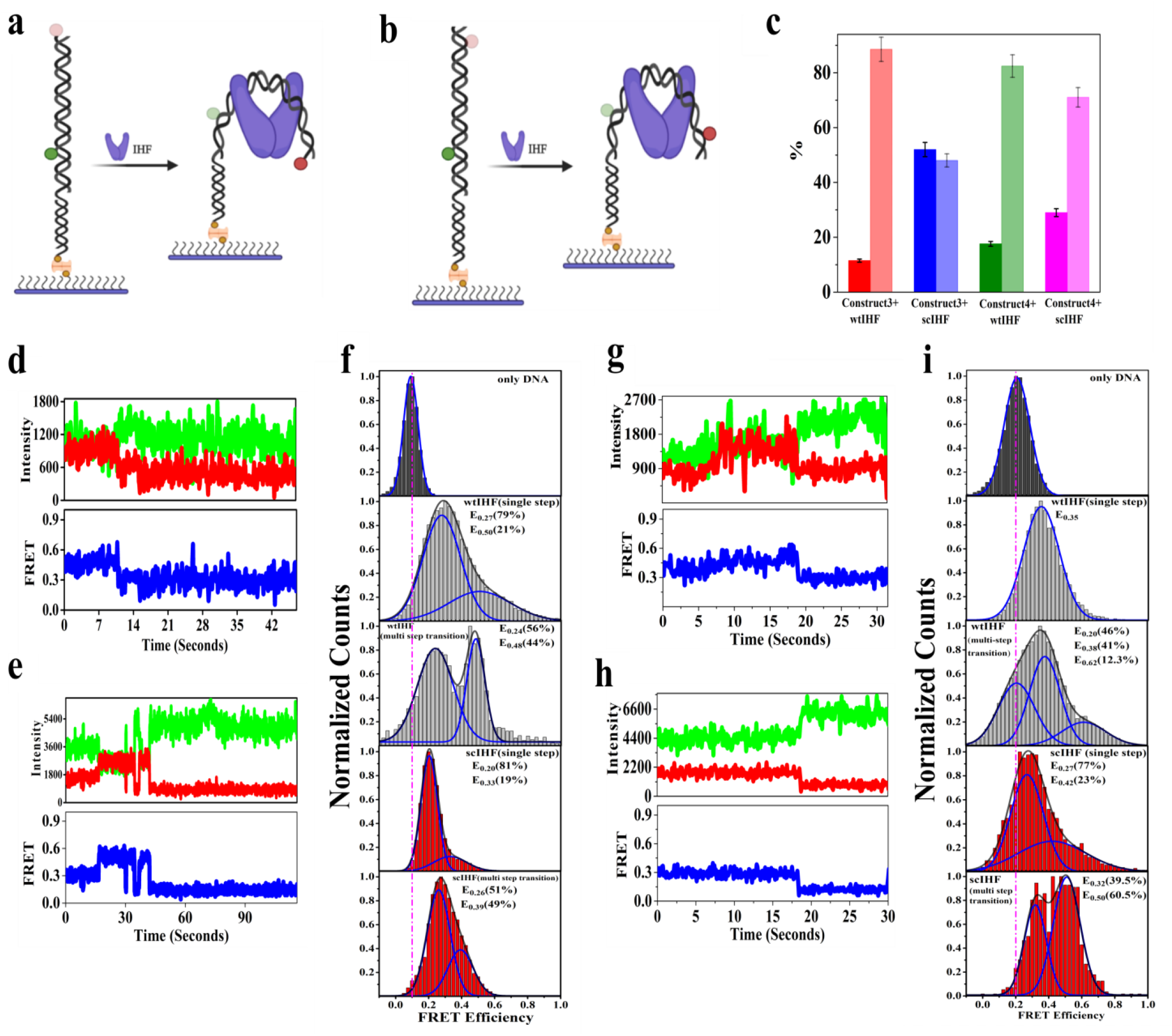
IHF induced bending of Construct 3 and 4 **(a)** and **(b)** Sketch of the experimental model with Construct 3 and 4 of DNA respectively. **(c)**The relative percentage of molecules in dynamic FRET states (dark-colored bars) and molecules existing in a constant static state (transparent bars). **(d)**and**(e)** Time traces of representative single-molecule and FRET efficiency plots of Construct 3 of DNA when allowed to bind with the proteins wtIHF and scIHF respectively. **(f)**Sequence of FRET Histograms illustrating only DNA Construct 3 followed by protein-bound (wtIHF and scIHF) DNA, molecules being separated based on their dynamicity. **(g)** and **(h)** FRET traces and corresponding efficiency plots of DNA Construct 4 with wtIHF and scIHF respectively. **(i)** FRET Histograms of Construct4 naked DNA, also with scIHF and wtIHF proteins.

**Figure 4.**
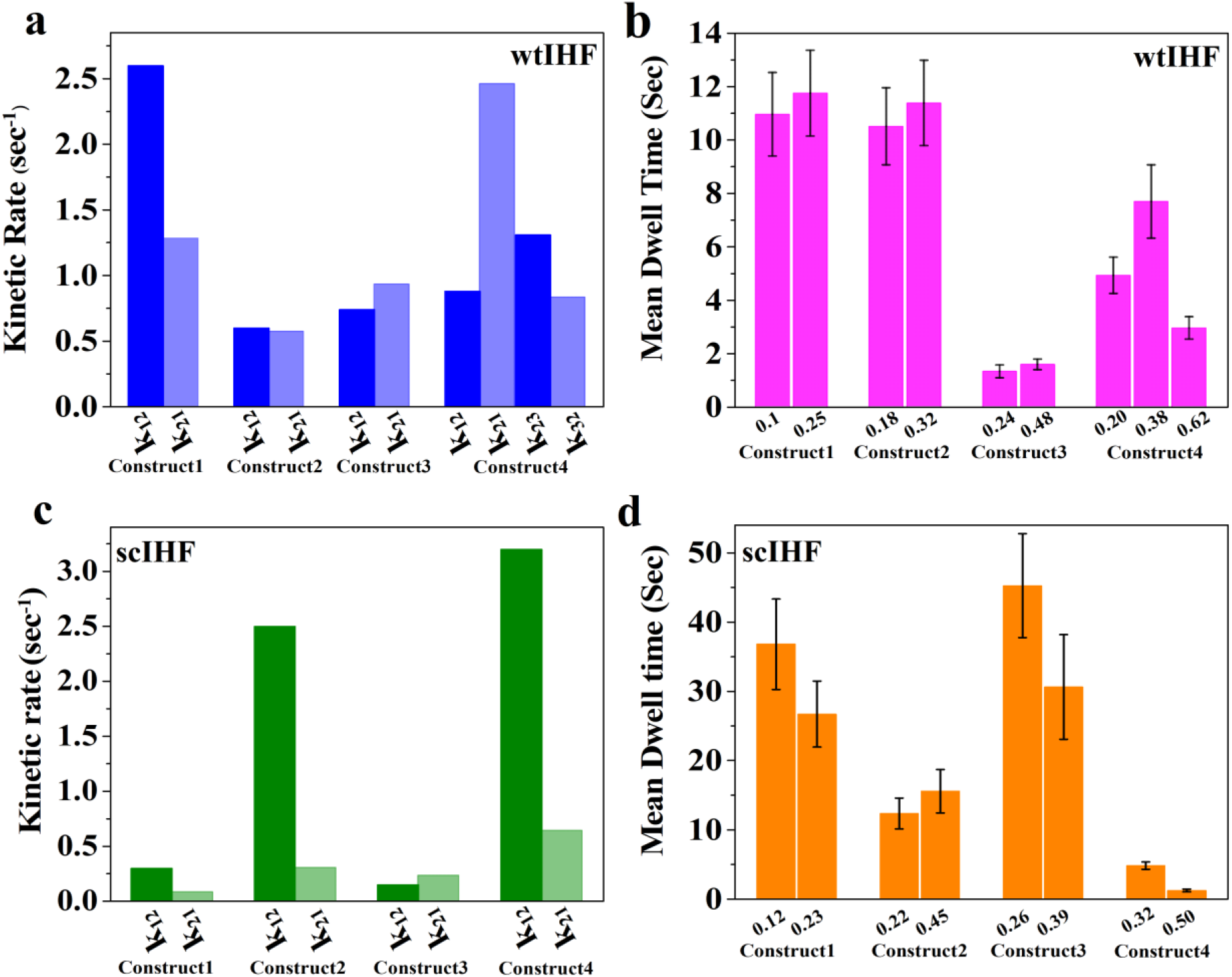
Comparison of kinetic rate transitions and dwell times between scIHF and wtIHF bound DNA **(a)** and **(c)** kinetic rates of transition from one state to next determined by applying HMM on molecules possessing multi-state behavior for all the four constructs of DNA when they are bound with wtIHF and scIHF respectively. **(b)** and **(d)** represents the mean dwell time at each particular FRET state as computed manually from individual time traces of the molecules exhibiting multi-state transition.

In order to explore clear information about the conformational rigidity of the protein-bound DNA by accumulating more information from the FRET changes between the straight and partially bent conformations, we altered the labelling strategy by shifting the acceptor position five nucleotides away from 5’ end, termed as Construct 2 (Figure 2b, Figure S1b, and Table S1). Here, the unbound DNA molecules demonstrated the maximum population at 0.1 FRET efficiency, the same as observed for the end-labelled construct. 21% of the scIHF bound DNA found to be in dynamic equilibrium between three conformational states, a highly populated state with *e*_*fret*_ around 0.45 along with two less populated states with *e*_*fret*_ around 0.22 and 0.7 (Figure 2c, Figure 2i). The mean dwell time at the low and high FRET region is found to be 12.3 seconds and 15.6 seconds respectively (Figure 4d). However, only 7% of the wtIHF bound DNA is found to show the dynamicity and exhibit interconversion between two FRET states, 0.18 and 0.32. Both of these low lying FRET states also have almost comparable mean dwell time values of 10.5 seconds and 11.4 seconds respectively (Figure 4b). Both of scIHF and wtIHF bound DNAs with a static trajectory till photo-bleaching show *e*_*fret*_ at∼0.25, with an additional less populated broad distribution peak at 0.4 for scIHF bound molecules (Figure 2i). Corroborating the results obtained considering the *e*_*fret*_ values and the population distributions and dwell time of each state suggests that the induced kink in the DNA by IHF does not acquire a uniform regular concave shape. Rather, the termini remain away from each other and the distance of the interior region from the end is more constricted. The kinetic rates of transition from and to each of the states attended during the dynamic transitions are shown in Figure 4. However, just like Construct 1, a sharp bending in the DNA is not observed for either of the proteins, as has been found previously by using ensemble FRET and other techniques.^40; 41^

### A-tract dynamics due to IHF induced bending

It has been well established that the process of dsDNA bending is not uniform in nature, the constriction and dilation of the arms of dsDNA continue differentially throughout the nucleic acid topology.^42; 43^ This is also an indicator of the heterogeneous nature of the specific complex formed between the DNA and bending protein. Relatively a broad distribution of bend angles has been previously reported from the atomic force microscopy measurements on some specific complexes that illustrate the fluctuations within a single free energy well in the conformational landscape of the complex. However, the resolution of the above studies are predominantly based on the single-peaked distributions of varying widths.^44^ With a view to get a profound perspective of the minute conformational and structural fluctuations, along with the mechanistic details of the interior of the dsDNA at A-tract region, the donor is labeled at the beginning of A-tract sequence from the binding site of the protein to decipher the dynamics of both the DNA constructs. In one of the constructs, the acceptor is labeled at the other end of the DNA (Construct 3, Supplementary Figure 1c) to gain a detailed perception of the A-tract vibration with respect to the end of DNA. In the other construct, the acceptor has shifted five nucleotides from the 5’end to monitor the interior constriction of the A tract region, termed as Construct 4 (Supplementary Figure 1d). While scrutinizing the conformational dynamics of the A-tract region compared to the end of DNA (Figure 3a), we found 52% of the molecules experience progressive alteration between two conformational states upon binding to scIHF (Figure 3c). The protein-bound DNAs were found to be shifted towards higher FRET state than those of the naked DNA molecules having maximum distribution at 0.1 FRET state, as observed for the previous two constructs (Supplementary Figure S2c, Figure 3f). The two conformational states of scIHF bound molecules fluctuate between *e*_*fret*_ values of 0.26 and 0.39 with almost equal population distribution (Figure 3f). The lower FRET state was more prolonged with a mean dwell time of 45.3 seconds than at the higher FRET state (30.6 s, Figure 4d). On the other hand, 12% of molecules attained dynamic conformational changes between two states while forming a complex with wtIHF, having peaks at 0.24 and 0.48 (Figure 3c, f). This could be due to the fact that wtIHF leads to a subtle more constriction of the A-tract zone with respect to the ends, compared to the bending by scIHF. Interestingly, the time spent at these two states is also comparable, *viz*. 1.3 seconds and 1.5 seconds (Figure 4b). The molecules staying in the same state until photo-bleaching showed an efficiency maximum at 0.2, with an additional small peak at 0.33 in the case of scIHF binding (Figure 3f). However, the scenario alters for binding with wtIHF, as the efficiency shows a peak shift towards 0.27 and 0.5 and the fraction of molecules exhibiting intermediate FRET state increases (Figure 3f), further reinforcing the evidence of an increased motile A-tract coming close to the DNA end while binding to wtIHF. Now, the study was continued shifting the acceptor position five nucleotides towards the interior of the DNA, Construct 4 (Figure 3b and Supplementary Figure S1d). The free DNA shows a populated peak at 0.2 (Supplementary Figure S2d, Figure 3i), indicating the FRET state of B-DNA conformation that primarily behaves like a stiff rod. After delivery of wtIHF protein and upon binding, about 18% of the molecules with conformational switching properties exhibit three distinct and well-populated peaks (Figure 3c), two being shifted towards higher FRET states (0.38 and 0.62) and one corresponding to the unbent state, 0.2 (Figure 3i). The mean dwell time is minimum (2.9 seconds) for the state with high FRET value, followed by the unbent conformation (4.9 seconds) (Figure 4b). The slightly constricted state has a maximum dwell time of 7.68 seconds (Figure 4b). However, the acquisition of FRET states due to the dynamics of DNA (29%) when exposed to scIHF exemplified two FRET states (0.32 and 0.5, Figure 3c), depicting the fact of capturing of a dilated bent DNA compared to the case observed in presence of wtIHF (Figure 3i). In presence of scIHF, the molecules appear shuttling between two noticeable conformations, with *e*_*fret*_ values 0.27 and 0.42, whilst upon wtIHF binding, there is an occurrence of one efficiency peak at 0.35 (Figure 3i). Compiling of the above results, it appears that the *e*_*fret*_ values for the local interim vibration of the 3’-end of the A-tract region with respect to the interior as well as the terminal end of the DNA in presence of both scIHF and wtIHF are higher than that of the constructs labelled at the terminal. This gives the indication that the internal region of the DNA closer to the induction of kinks by protein is more constricted than that of the terminal ends of the DNA, suggesting induction of an irregular semi-circular bend by the DNA.

However, the bending in the DNA captured in the crystal structure due to binding with both scIHF and wtIHF is more compact and constricted (Supplementary Figure S3 and Table S2).^6; 7^ Our results from the smFRET experiments show a more loosely bent form of the DNA or rather the relaxed nature of the complex. This observation directs towards two important insights. First, the clamping of the proteins causes either of the arms of the DNA to bend increasing the distance between both of the arms and hence, the value of *e*_*fret*_ decreases, second, it is also possible for both the arms bend in a wobblier fashion, resulting in multiple dynamic states. The utility of the constricted crystal state has been taken forward to measure the angle of flexibility where the set of maximum supporting angular values have been exemplified from the crystal structure value.

### A quantitative study of wtIHF induced DNA compaction

The employment of smFRET has immensely helped in quantifying the conformational switching of the nucleoprotein complex at nanoscale resolution. As the proteins induce two kinks in the DNA, we have quantitatively interpolated the compaction of the dsDNA through the calculation of the flexure angle, considering the outline of the bent DNA as a quadrilateral. Corresponding to each of the FRET Efficiency values, a set of angular values is obtained having a distribution range of maximum 8-9 degrees. For Construct 1, the angle of flexibility was determined for the two sets of DNA bend conformation. The average equivalent of angle for 0.22 FRET Efficiency was found to be 91.7 ± 1.1 degrees (Mean ± SD.), with a maximum distribution of 90.8 degrees. The other conformation with FRET Efficiency 0.25, the angular bend came out as an average of 89.0 ± 1.1, with a maximum distribution at 88.5 degrees (Supplementary Table S4a and Figure 5a). Furthermore, we also have attempted to assign an erratic degree of plasticity to the DNA-protein complex at different sites of the dsDNA by labelling it at different interior positions, as explained in the previous section. In Construct 2, the DNA in the bent state attains three different conformational states. And, the mean angles of flexibility for the three different states are 105.8 ± 0.5, 97.9 ± 0.3 and 92.1 ± 0.2 degrees respectively (Supplementary Table S4b and Figure 5b). The ranges of angular values, obtained are found to be quite constricted between 1-2 degrees for this DNA construct. In previous studies, the bend-angles, calculated as the supplementary to the angle of intersection between two bent arms of DNA considering them as tangents, indicates that IHF induces approximately 60-80 degrees bending in the DNA.^45^ This result from the AFM measurement is close to the angle of flexibility while monitoring the end-labelled constructs in our study. However, the minuscule level of analysis due to the introduction of two kinks by the protein remained unaddressed through AFM imaging. Also, investigation of the curvature of the dsDNA with respect to the A-tract region was not done in the AFM study. Analysis of the extent of bending of DNA by several other proteins has also been explored by AFM, among them DNA Pol β, MutS, and Rad4-Rad23 known to induce light bends in the DNA whereas IHF, Topoisomerase II, and HMGB1 Box introduce sharp kink in the DNA.^45; 46^ In Construct 3, the donor has shifted ten nucleotides towards the interior in the region of the A-tract and four states of DNA bent confirmations are identified. The flexure angle is expanded to 121.23 ±1.80 degrees for the lowest FRET state of the DNA, as attains the most dilated form. The flexure angle eventually decreased with the compressed states of the DNA upon binding to the protein and found to be 117 ±1.9, 100.3± 2.2, and 99.0 ± 2.2 degrees (Supplementary Table S4c and Figure 5c) analogous to the increasing FRET Efficiency. In Construct 4, of interiorly labelled both donor and acceptor and as a consequence, the length of the arms of the quadrilateral decreased, giving to an augmented flexure angle value. Thus, even though the distance between the two arms of the bent DNA while complexed with IHF is less, the evaluated values of flexure angles are elevated in the case of Construct 4. The three different bent conformations of DNA give rise to angular values as, 127.1 ± 1.7, 124.2 ± 1.8, and 103.4 ± 0.5 degrees (Supplementary Table S4d and Figure 5d). Computation of the angles with respect to different sections of DNA provided us the information that the angular values increase as we move towards the interior of the DNA. This gives a model of more protuberance in the flexibility of DNA in the interior region close to the protein-bound site than at the ends.

**Figure 5.**
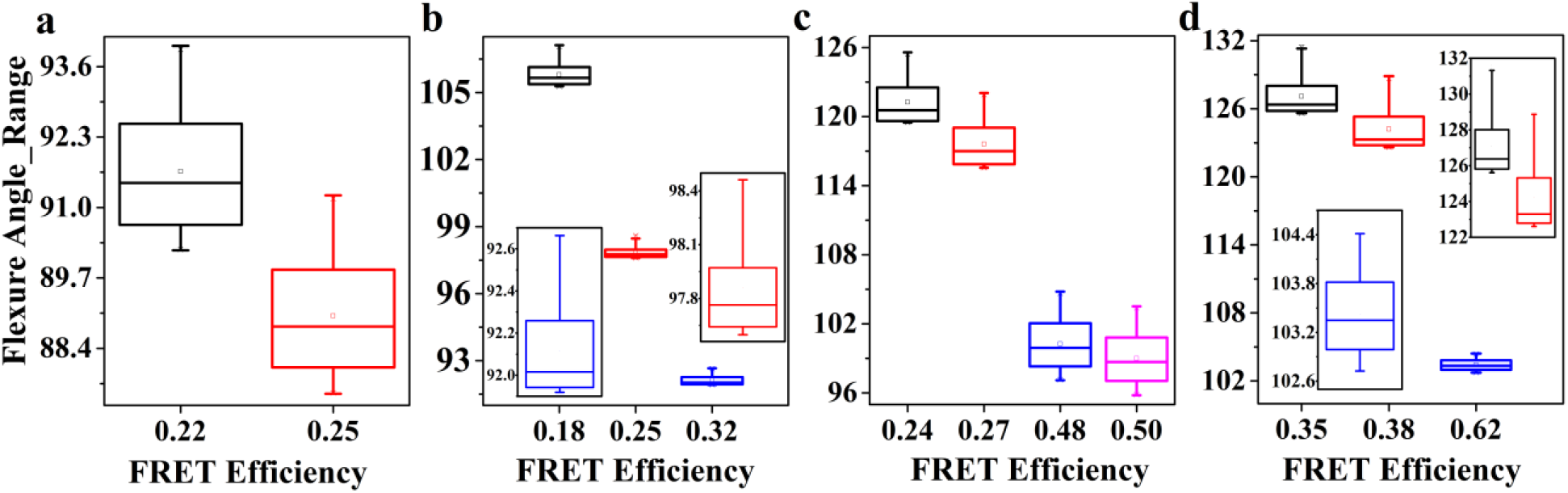
Quantification of flexure angles upon wtIHF binding. Boxplots of the distribution of flexure angle values (in degrees) at each FRET states depicting the bent population of DNA when it is bound with wtIHF, **(a)** to **(d)** represents the sequence of Constructs1 to 4. The narrow distribution of the angles leads to constricted box-plots, as are expanded as an inset of the figure.

### Comparative bending analysis by scIHF

We have also characterized the dynamics of dsDNA upon binding to scIHF binding through the additional insight of quantification of the degree of flexibility of DNA. The lesser the value of the angle, the sharper is the curve ensuring tight compaction, thus the degree of flexibility is the route towards quantification for the acquirement of compactness. Here, we have exemplified that switching between multiple dynamic states, from a compact genome to a loose dilated one in protein-bound condition changes the extent of bendability of the DNA in the same DNA-protein complex. The DNA acquires three bent states in Construct 1, and the calculated mean values of flexure angles are 102.1 ± 1.7, 98.1 ± 1.8, and 91.8 ± 1.3 degrees (Supplementary Table S5a and Figure 6a) and found to be decreasing with the increase in *e*_*fret*_ values. Similarly, four distinct structural forms are identified in Construct 2. The maximum bending angle found is 99.21 ± 0.35 degrees, corresponding to the highest FRET state (Supplementary Table S5b and Figure 6b). The angular value eventually decreased to 97.1 ± 0.3, 82.02 ± 0.2 and 69.6 ± 0.1 degrees with the increase in compactness of the genome (Supplementary Table S5b and Figure 6b). The attainment of the maximum distribution of the angles in all the individual cases has been depicted in Supplementary Figure S8 by extrapolating the values of angles in the form of a histogram. Shifting the donor fluorophore towards the A-tract region, just as we found in the case of wtIHF binding, the angles increased compared to the Constructs 1 and 2 even in the range of close FRET states. The most constricted conformer exhibited the angular value of 108.4 ± 2.1 degrees which is larger than the maximum angular values found in the Construct 1 and 2. The eventual dilated states led to increment in the flexure angle values, i.*e*. 113.6 + 2.0, 120.9 + 1.8 and 129.6 + 1.5 degrees (Supplementary Table S5c and Figure 6c). The flexure angles are also determined for Construct 4, the angles of the flexibility of DNA bent state has been found to range from 136.2 ± 1.5 to 112.0 ± 1.0 degrees (Supplementary Table S5d and Figure 6d) following the FRET Efficiency values from low to high FRET states. The flexure angles are larger for the Constructs 1, 3, and 4 while they form a complex with scIHF compared to wtIHF, whereas the same is higher for Construct 2 wtIHF complex. The standard deviation (SD) of the values of bending angles obtained has been plotted for each of the constructs with the proteins, wtIHF, and scIHF. It is worth mentioning here that the values obtained from the smFRET study are comparatively lower compared to the same reported earlier by using other methods.^46^ The pattern of SD values for all sets of DNA constructs is found to be close, except for Construct 4 where the distribution of angles analogous to the maximum FRET state illustrates a drop in the SD value (Figure 7).

**Figure 6.**
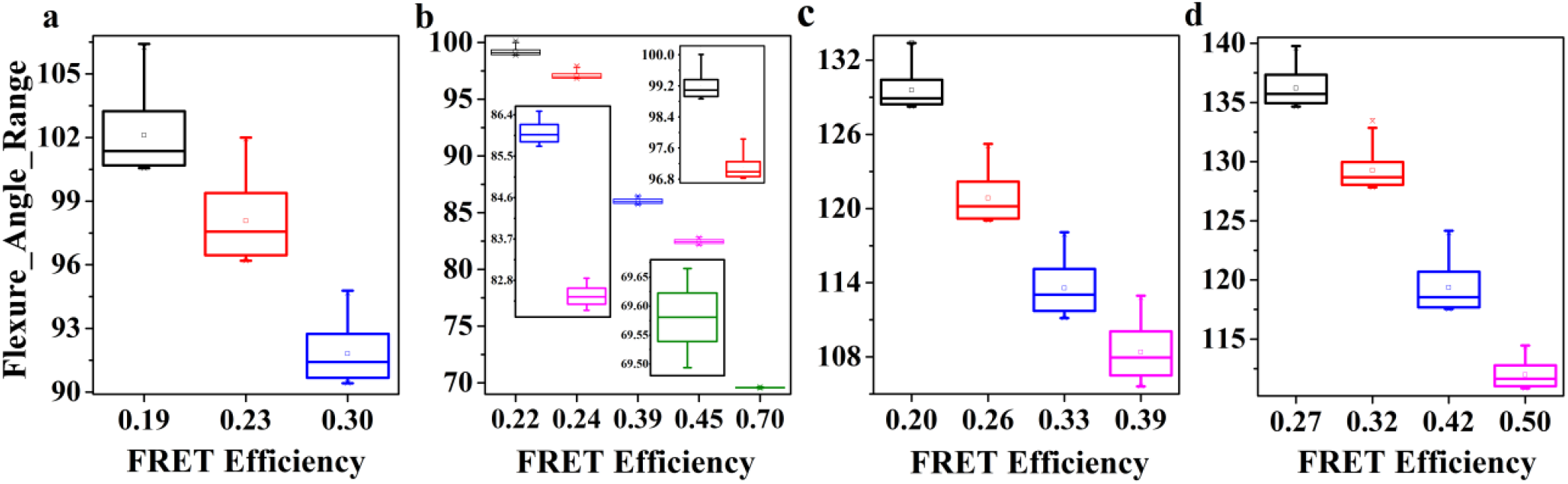
Quantification of flexure angles upon scIHF binding. Boxplots of the distribution of flexure angles, when DNA is bound with scIHF. The x-axis represents the corresponding bent FRET state and the y-axis covers the distribution of the range of angles. Just as the previous figure, **(a)** to **(d)** stands for construct1 to 4 in the order.

**Figure 7.**
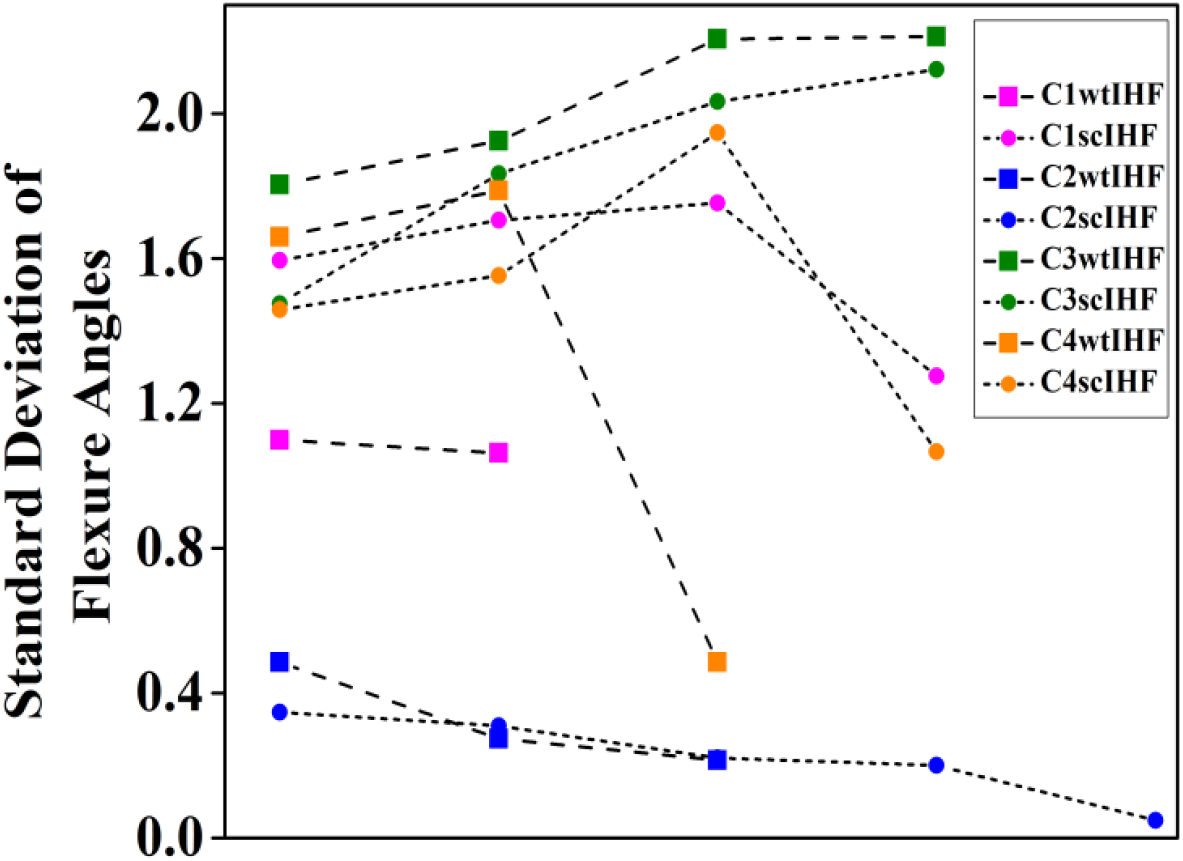
Variation in flexure angles between scIHF and wtIHF induced DNA bending. Correlation of Standard deviation of the range of flexure angle values corresponding to all the set of FRET states for each constructs when wtIHF and scIHF bind with the DNA, (left to right indicate increasing order of FRET Efficiency values). The circles represent scIHF and boxes represent values corresponding to wtIHF. The similar color goes for a similar construct of DNA.

## Discussion

Life has diversified in the most spectacular ways during the past 3.8 billion years, evolving into Bacteria, Archaea, and Eukaryota^47^ from a small population of unicellular entities *i*.*e*. Last Universal Common Ancestor (LUCA). ^48; 49; 50^ Astonishingly, underneath this remarkable diversity from molecular to organism level, one particular theme that nature has conserved across all the domains of life is the ability to store information in nucleic acids (DNA or RNA). It also ensures its existence and propagation through generations employing precise and highly regulatory compaction mechanisms. The 1.5 millimetres long *E*.*coli* genome consists of approximately 4.6×10^6^ base pairs. The intrinsic bends on the DNA curvature due to Brownian motion along with regulated bends by the NAPs make it possible to accommodate the DNA within a 3 µm long × 1µm diameter cell.^51^ Out of many known NAPs, we have quantified the IHF induced bending in this study by taking a dsDNA shorter than the persistence length (50 nm or 150 base pairs), thereby ruling out the possibilities of having any intrinsic bends.^52^ Whether the bending of DNA is induced or intrinsic can be categorized into two major models, namely, ‘Junction Model’^53^ and ‘Wedge Model’.^54^ The concept of wedging is used to describe the bending of DNA, if small wedges are inserted at a particular nucleotide region and the final curvature of DNA is the sum of all individual wedges. However, the junction model illustrates the introduction of a kink between two consecutive nucleotides and the DNA helix returns to the canonical B-form structure just after the bend. Hence, bending by IHF appears to fall more accurately into the junction model that is also evident in the previously reported crystal structure analysis.^6^

It has been reported earlier that binding of IHF with the DNA produces two kinks, one at the consensus sequence (H’ sequence) and the other close to it with a gap of nine base pairs between them.^6^ In order to provide a realistic illustration in agreement with the existing crystal structures, a ‘C’ shaped bending was assumed in this study. Although the previous structural investigation on IHF suggested that it promotes sharp bending on the overall DNA contour that freezes the nucleoprotein complex in one particular conformation^6; 7^, the likelihoods of the existence of multiple bend conformational states could not be ruled out. Also, in a physiological system, it is undeniably plausible that the behaviour of every individual biomolecule is heterogeneous in nature even if they are present in a similar immediate microenvironment. The previous attempts on capturing the dynamics of IHF-DNA have reported representation of a partially bend state by only 25-30% population whereas the rest mimics the sharp bend (larger than 160 degrees) state, as captured in the crystal structures.^55^ Surprisingly, we did not observe any bend state that mimics the crystal structures rather, the majority of the population across different experimental conditions exhibit a partially bend conformation. There could be three possible explanations for this ambiguity observed. Firstly, unlike our continuous stretch of dsDNA, the dsDNA construct used in the crystal structure had a nick near the A-tract region and close to the kink site to facilitate the crystal packing^6^ and this might have helped IHF to bend efficiently as IHF bending efficiency has been reported to be sequence-dependent.^55^ Secondly, in our experimental approach, one end of the dsDNA was immobilized on the slide surface. This might have hindered the induction of sharp bends. Thirdly and most importantly, the experimental time frame of the two approaches is drastically different, as probing of conformational dynamics by smFRET occurs in millisecond resolution whereas formation of a crystal is a matter of several hours to days, that might induce a sharp bend even with a continuous stretch of dsDNA. We have considered a two dimensional (2D) projection of the entire three dimensional (3D) reality and determined the flexure angles as it defines the apparent bending of DNA more explicitly and to provide a realistic picture of the degree of flexibility taking into account of the local reduction of stiffness (Supplementary Figure S4 and Figure S5).^56^

By thoroughly evaluating the events of conformational heterogeneity in the course of appraising molecule by molecule, we confirm that the IHF-DNA nucleoprotein complex adopts an ensemble of ‘slacked-dynamic’ conformations (Figure 8 and Figure 9). From the experimental observations and analysis, we report a differential population distribution in the context of transition between different conformational states for the two variants of IHF in the same experimental conditions. Without any exception for all sets of experiments, the transient state population is found to be higher for scIHF. However, it has not escaped our careful observation that the ‘conformational window’ i.e. the range of different conformers was slightly broader despite having a smaller transient population in the case of wtIHF. Also, we have observed a confirmation by non-dynamic molecules that lie outside our transient ‘conformational window’ in three of the constructs, out of four dsDNA constructs used for the study. This could be due to the fact that most of the time wtIHF is able to trap the DNA in the most energetically favourable conformation and as a result of that, we see a lower transient population. This also suggests that although the scIHF is more efficient in DNA bending in a biophysical context in terms of the introduction of flexibility, perhaps wtIHF is even more efficient at it in a ‘biology’ context. This behaviour of the wild-type variant might have important evolutionary consequences in gene regulation and genome organization. In eukaryotes, the chromosome is composed of both nucleosome and non-nucleosomal histone-DNA complexes known as ‘pre-nucleosomes’, that are later converted into higher-ordered canonical nucleosome structures by several chromatin remodelling proteins.^57^ The wide range of the calculated flexure angles forms our experiments suggests that IHF when bound to the DNA, acts as a pre-nucleoid structure that can be thought of as a ‘nucleation point’ for higher-order genome organization in prokaryotes. The pre-nucleoid structure further requires organization by other NAPs like heat-stable protein from *E*.*coil* strain U93 (HU), histone-like nucleoid structuring protein (H-NS), and structural maintenance of chromosome (SMC) proteins, those ensure higher-order compaction. This hypothesis can be further supported by the fact that HU preferentially binds to a bend DNA^58^ and HNS acts as a ‘bridge’ between two different DNA filaments that forms a loop (‘Daisy Chain Model’). ^23; 59; 60^ SMC proteins are also well known for their loop forming abilities that have been appealingly described through ‘One ring, two DNA Model’ and ‘Hand-cuffed Model’^61^. Following our observations, this dynamic or transient behaviour of biomolecules has been discovered in many other systems as well, ranging from viral proteins to eukaryotic membrane proteins and nucleosomes. ^33; 34; 62^ The ability of biomolecules to acquire different conformations have been proved to play crucial roles in enzyme promiscuity, catalysis, substrate binding, and product release, and hence, directly associated with its capability to evolve. ^63^

**Figure 8.**
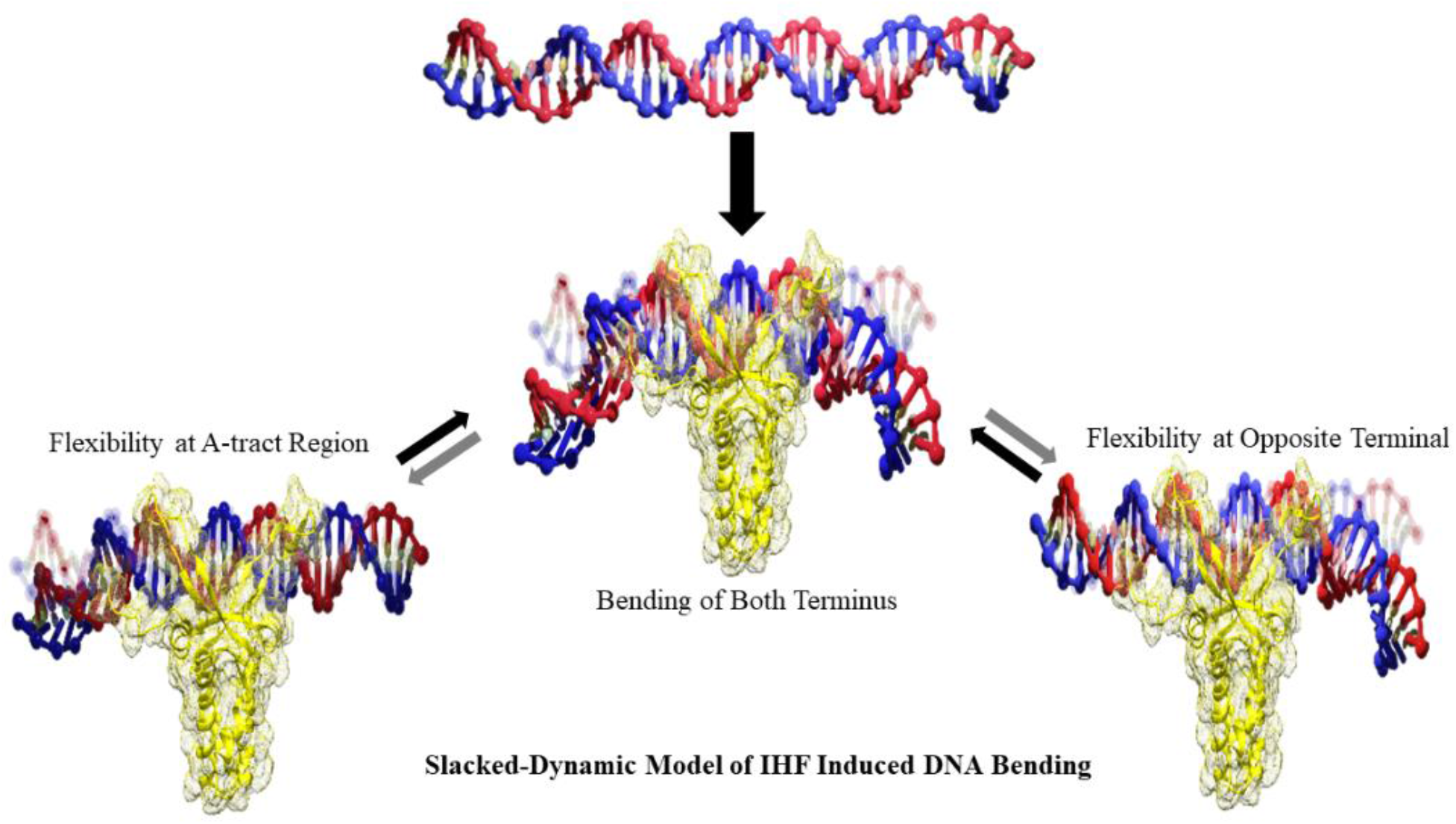
Final model of bending of the dsDNA by IHF. The model depicts the dynamic transitions between the conformations. The models were made with oxDNA^75^ and Blender^76^.

**Figure 9.**
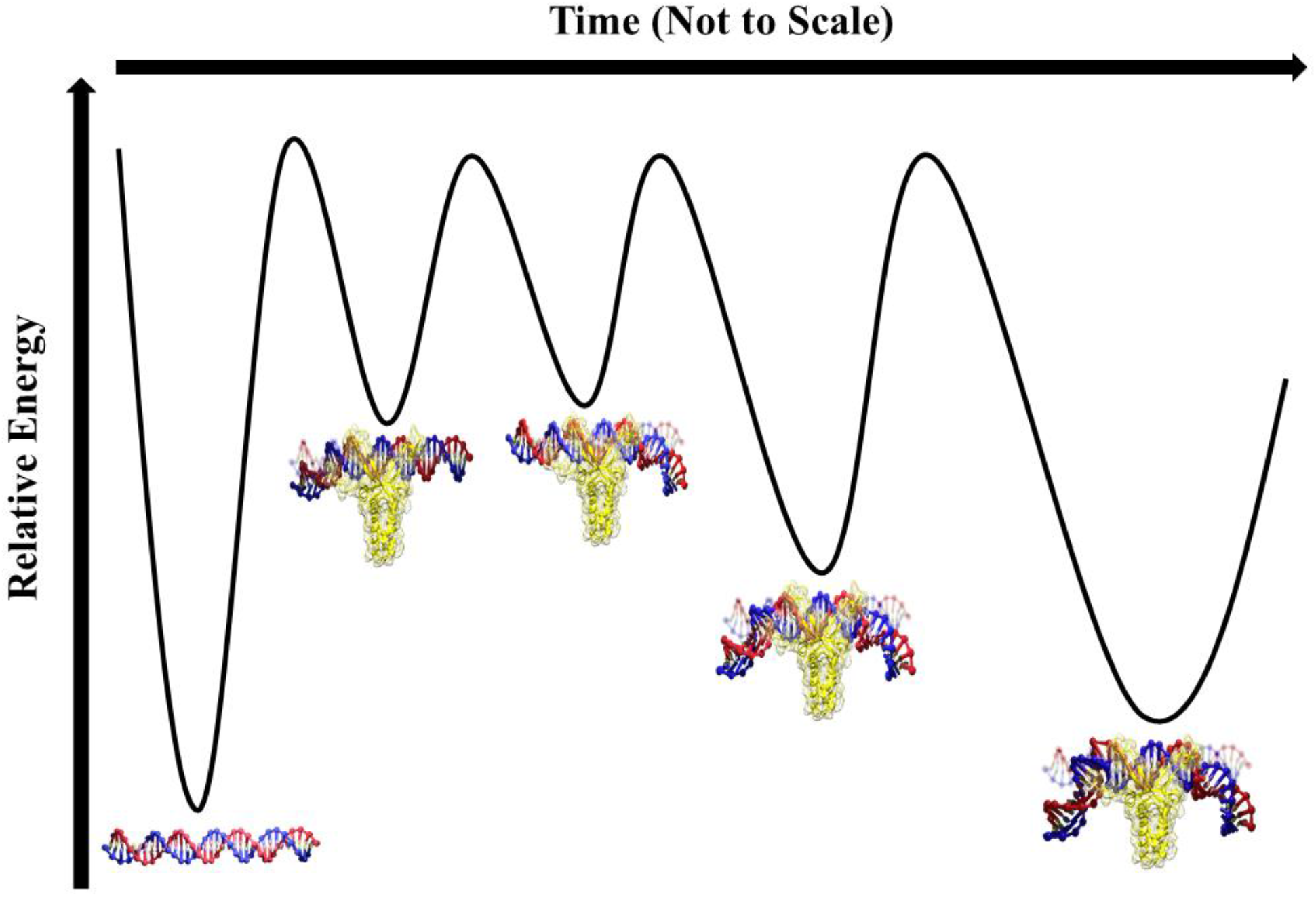
Schematic representation of free energy landscape of the IHF-DNA complex. The conformations predicted from our experimental observations (Figure 8) are represented here in terms of their relative stability. A hypothetical more stable conformation which closely resembles the crystal structure has been depicted here as well.

Additionally, our findings provide crucial insights regarding the possible role of IHF as a transcription regulator in bacterial cells. In continuation with the argument and facts we have presented in the earlier section, several previous studies also have established a link between the transcriptional activity and the number and stability of loops at a specific DNA region.^64; 65; 66^ We hypothesize that the wobbling states of the IHF-DNA nucleoprotein complex impart a certain degree of plasticity in the promoter regions that in turn leads to the recruitment of not only RNA Polymerase but also several other regulatory proteins including σ^54^, a transcription factor at the enhancer region as has been verified in the previous studies.^67^ The dynamic topology of the semi-looped DNA could physically disrupt the interactions among the DNA bound transcription factors, thereby suppressing the transcription or can promote cooperative binding by providing room for other proteins by continuing their respective function without being sterically affected. Another illustration concerning the flexibility of the dsDNA even in packaged form can be taken forward to the DNA repair mechanism. A recent study demonstrated that chromatin acts as a better template than naked DNA in homology-directed DNA repair.^68^ Achievement of dynamicity upon being bend by architectural proteins makes the DNA a better target to enter into the DNA repair pathway.^69^ Thus, there is a high possibility that in the case of bacterial nucleoids and repair proteins, directly or indirectly, NAPs may involve in a plethora of molecular pathways in the cell and their exact functions in those are beyond our understanding.

In summary, we would like to emphasize that our study provides a detailed description of chronological events during the bending of a dsDNA by the two IHF variants by analyzing the FRET profile at single-molecule resolution. This not only allows the scope of comparison but also opens up an opportunity to understand the functions of other nucleoid-associated proteins (NAPs) in general. Apart from satisfying the scientific curiosity, these insights might find its applications in the pharmaceutical industry as IHF has been identified as a key factor that determines the overall integrity of biofilms in several deadly human pathogens.^24,25^Finally, we hope that these findings regarding the conformational dynamics or flexibility of the IHF-DNA nucleoprotein complex at a millisecond time scale will help to understand the biology better by complementing with the previous detailed atomic-level structural analysis, that eventually will allow accessibility towards better hypothesis and understanding about the NAPs.

## Materials and Methods

All the chemical reagents used are of analytical grade and were purchased from Sigma-Aldrich and Merck. All the common salts used for buffer preparations and bacterial culture media were purchased from Sisco Research Laboratories Pvt. Ltd. (SRL). The media for cell culture are from HiMedia Laboratories.

### Induced Over-Expression and Purification of single-chain Integration Host Factor (scIHF) and wild-type Integration Host Factor (wtIHF)

scIHF gene was cloned in plasmid pETscIHF having Ampicillin antibiotic-resistant marker, and the plasmid containing the wtIHF gene was a generous gift from Prof. Peter Droge of Nanyan Technological University.^70^ For induction and protein over-expression, plasmid pETscIHF and the plasmid with wtIHF were transformed into *Escherichia coli* BL21DE3 cells. 2 liters of Luria-Bertani broth media supplemented with Ampicillin (100 µg/ml) was prepared and was inoculated freshly with over-night grown cultures of transformed *Escherichia coli* BL21DE3 cells. The cells were grown at 37°C for approximately 2.5 hours until the absorbance of the culture at 600 nm reached 0.4. At this point, induction was initiated by adding 2 mM of Isopropyl β-D-1-thiogalactopyranoside (IPTG) into the culture media. The cultures were grown at 20°C for 12 hours after induction. Then the culture media were centrifuged (Heraeus, Biofuge, Stratos, Thermo Fisher Scientific) at 4000 rpm at 4°Cfor 15 minutes, and the cells were collected.

Cells were then re-suspended in Lysis Buffer (20 mM Tris-Cl, pH 8.0, 20 mM NaCl, 1 mM EDTA, 1 mM PMSF, 10% Glycerol) and then sonicated (Ultrasonic Processor, Cole Parmer) using 5 cycles of 30 seconds ‘ON’ and 30 seconds ‘OFF’ sonic pulses with 14% amplitude. The cell lysate was centrifuged (Heraeus, Biofuge, Stratos, Thermo Fisher Scientific) at 10,000 rpm at 4°C for 1 hour, and the supernatant was collected. The supernatant was subsequently loaded manually in a pre-equilibrated Ni-NTA affinity chromatography column and allowed to pass through the column with a flow rate of 0.5 ml/min. The column was then washed thoroughly with 5 CV (column volume) of Wash Buffer (20 mM Tris-Cl, pH 8.0, 20 mM NaCl, 1 mM EDTA, 40 mM Imidazole) keeping the same flow rate. The elution was done with 10 ml of Elution Buffer (20 mM Tris-Cl, pH 8.0, 20 mM NaCl, 1 mM EDTA, 400 mM Imidazole) with a flow rate of 0.5ml/min. The presence and purity of the protein are checked by Bradford Assay followed by Polyacrylamide Gel Electrophoresis (SDS-PAGE, 5% Stacking, and 12% Resolving). The proteins were then concentrated in a Storage Buffer (25 mM Tris-Cl, pH 8.0, 25 mM NaCl, 25% Glycerol) using 10 kDa Amicon Ultra-centrifugal Filters (Merck). The concentrated scIHF or wtIHF were collected in micro-centrifuge tubes in small aliquots (100 µl), fast-freezed, and stored at -80°C till further use. The entire purification process was performed at 4°C.

### Design of DNA Constructs

Custom made HPLC purified fluorescent labelled or unlabelled DNA oligonucleotides were purchased from Integrated DNA Technologies, Inc (IDT). The oligonucleotides were designed keeping in mind the specific binding site for single-chain (sc) and wild-type (wt) variants of Integration Host Factor (IHF). Four different combinations of DNA constructs were made keeping the nucleotide sequence the same and changing the position of the fluorescence probes (Cyanine 3 as Donor and Cyanine 5 as acceptor). A short biotinylated (at 5’ end) oligonucleotide sequence was designed for the surface immobilization of the DNA constructs. The sequences and point of labelling have been mentioned in detail in the supplementary information (Supplementary Figure S1 and Table S1).

To carry out the experiments, appropriate sequence pairs were selected to generate a FRET pair, along with the biotinylated sequence (Biotin:Cy3:Cy5=1:1:1) in T50 Buffer (10 mM Tris-Cl, pH 8.0 and 50 mM NaCl). The final concentration was kept at 20µM for each of the sequences. Annealing was done by heating the mixture at 95°C and then gradually cooling it down to 8°C. The final experimental ‘single-molecule concentration’ (pico-molar level) was achieved by further serial dilution of the annealed product in Reaction Buffer (20 mM Tris-Cl, pH 8.0, and 10 mM MgCl_2_). The remaining annealed product was stored at -20°C for future use.

### smFRET Experiments and Data Acquisition

Pre-drilled quartz slides were processed to immobilize the DNA of interest as has been explained in our previous study.^38^ For immobilization, Streptavidin (0.2 mg/ml) was flown to the sample-chamber followed by washing and flushing with pre-annealed fluorescent labelled DNA with a biotin tag. Once there are approximate 400 molecules on the Cy5 channel of the emCCD camera (Andor ixon3 897), the unbound DNA samples were washed using the Imaging Buffer (Reaction Buffer supplemented along with 1 mM Trolox, 1 µg/ml Glucose, 1 mg/ml Glucose oxidase, and 0.03 mg/ml Catalase).^71^ At this stage, 500 nM of purified scIHF or wtIHF (in Imaging Buffer) was introduced inside the micro-fluidic chambers and data acquisition was initiated.

All smFRET experiments were performed using our home-built Prism-type Total Internal Reflection Microscopy setup.^72^ The proper focus over the samples was achieved with the help of an inverted fluorescent microscope (Olympus IX 71). Fluorescence signals from both Cyanine 3 and Cyanine 5 were collected using water immersion 60X, 1.2NA objective (Olympus). The signals were passed through a 550nm long-pass filter (Chroma) present inside the filter cube of the microscope and then Cyanine 3 and Cyanine 5 emission signals were separated using a 640nm DCXR (Chroma). Finally, the Donor and Acceptor emissions were projected separately onto the one-half of the Electron-Multiplied-Charged-Coupled Device Camera (EMCCD, Ixon3+897, Andor Technologies). The movies were recorded keeping a Multiplication Gain of 150 and Frame Integration Time of 35 or 100 milliseconds, using Visual C++ based data acquisition software from Prof. T. J. Ha.^73^ All the experiments and data acquisitions were performed at room temperature (25°C).

### Data Analysis

Raw movie files were analyzed using codes written on the Interactive Data Language (IDL) platform, a generous gift from Prof. Tae-Hee Lee of Pennsylvania State University.^74^ Time traces of individual molecules were analyzed and molecules having anti-correlated donor and acceptor signal with single-step photo-bleaching were selected. The molecules are separated based on the fact that whether they acquire multiple dynamic states or dwell in a single state till photo-bleaching and separate histograms are generated for each set of molecules. To determine the kinetic rate of transition from one state to another, we have employed the Hidden Markov Model Analysis. The kinetic rate of transition from one state to another is acquired by dividing the transition matrix outputs of the HMM codes by exposure time of the camera, in this case, 100 or 35 milliseconds. To extract the dwell time at a certain state, the time of stay at that particular state of several molecules showing multiple state changes were determined and the mean value has been reported here as the dwell time in the exact state.

### Calculation of Flexure Angles

The methodology for the estimation of the flexure angle has been described in detail in the supporting information (Supplementary Figure S4, Figure S5, and Table S2).To get a comparison of the range of flexure angle values for a particular DNA construct and Protein, the range of distribution of the flexure angles corresponding to every E_FRET_ value for each set of experiments are represented in boxplot (Figure 5 and Figure 6). A 3-dimensional plot of α, β and corresponding flexure angles are generated and also based on the number of times of occurrence of a particular flexure angle value for the range of α and β values in a particular DNA construct, a histogram is generated to get the details of the angular value having maximum distribution (Supplementary Figure S8, Figure S9, and Table S3).The codes for flexure angle calculations were written in Mathematica and all the plots are constructed using Origin 9.0.

## Supporting information

IHF_Supplementary

## Acknowledgments

The authors would like to thank Dr.Arghya Mukherjee and Debanjan Bandyopadhyay for their help in the calculations of flexure angle; Dr.Oishee Chakrabarti and Dr. H. Raghuraman for their valuable scientific inputs.

## References

1. Frishman, D. & Mewes, H.-W. (1997). PEDANTic genome analysis. Trends in Genetics 10, 415–416.

2. Luscombe, N. M., Austin, S. E., Berman, H. M. & Thornton, J. M. (2000). An overview of the structures of protein-DNA complexes. Genome biology 1, reviews001.

3. Torres-Escobar, A., Juárez-Rodríguez, M. D. & Demuth, D. R. (2014). Integration host factor is required for replication of pYGK-derived plasmids in Aggregatibacter actinomycetemcomitans. FEMS microbiology letters 357, 184–194.

4. Verma, S. C., Qian, Z. & Adhya, S. L. (2019). Architecture of the Escherichia coli nucleoid. PLoS genetics 15, e1008456.

5. Hołówka, J. & Zakrzewska-Czerwńska, J. (2020). Nucleoid Associated Proteins: The Small Organizers That Help to Cope With Stress. Frontiers in Microbiology 11, 590.

6. Rice, P. A., Yang, S.-w., Mizuuchi, K. & Nash, H. A. (1996). Crystal structure of an IHF-DNA complex: a protein-induced DNA U-turn. Cell 87, 1295–1306.

7. Bao, Q., Chen, H., Liu, Y., Yan, J., Dröge, P. & Davey, C. A. (2007). A divalent metal-mediated switch controlling protein-induced DNA bending. Journal of molecular biology 367, 731–740.

8. Robertson, C. A. & Nash, H. A. (1988). Bending of the bacteriophage lambda attachment site by Escherichia coli integration host factor. Journal of Biological Chemistry 263, 3554–3557.

9. Thompson, J., Fong, H. & Mark, L. (1988). Functional and structural characterization of stable DNA curvature in lambda attP. Structure and expression: proceedings of the Fifth Conversation in the Discipline Biomolecular Stereodynamics held at the State University of New York at Albany, June 2-6, 1987/edited by MH Sarma & RH Sarma.

10. Stojkova, P., Spidlova, P. & Stulik, J. (2019). Nucleoid-associated protein HU: A lilliputian in gene regulation of bacterial virulence. Frontiers in cellular and infection microbiology 9, 159.

11. Kamashev, D. & Rouviere-Yaniv, J. (2000). The histone-like protein HU binds specifically to DNA recombination and repair intermediates. The EMBO journal 19, 6527–6535.

12. Mikkel, K., Tagel, M., Ukkivi, K., Ilves, H. & Kivisaar, M. (2020). Integration host factor IHF facilitates homologous recombination and mutagenic processes in Pseudomonas putida. DNA repair 85, 102745.

13. Zhou, H.-X. (2011). Rapid search for specific sites on DNA through conformational switch of nonspecifically bound proteins. Proceedings of the National Academy of Sciences 108, 8651–8656.

14. Dhavan, G. M., Crothers, D. M., Chance, M. R. & Brenowitz, M. (2002). Concerted binding and bending of DNA by Eschericia coli integration host factor. Journal of molecular biology 315, 1027–1037.

15. Slutsky, M. & Mirny, L. A. (2004). Kinetics of protein-DNA interaction: facilitated target location in sequence-dependent potential. Biophysical journal 87, 4021–4035.

16. Kuznetsov, S. V., Sugimura, S., Vivas, P., Crothers, D. M. & Ansari, A. (2006). Direct observation of DNA bending/unbending kinetics in complex with DNA-bending protein IHF. Proceedings of the National Academy of Sciences 103, 18515–18520.

17. Vivas, P., Velmurugu, Y., Kuznetsov, S. V., Rice, P. A. & Ansari, A. (2012). Mapping the transition state for DNA bending by IHF. Journal of molecular biology 418, 300–315.

18. Ellenberger, T. & Landy, A. (1997). A good turn for DNA: the structure of integration host factor bound to DNA. Structure 5, 153–157.

19. Travers, A. (1997). DNA–protein interactions: IHF-the master bender. Current Biology 7, R252–R254.

20. Biek, D. & Cohen, S. (1989). Involvement of integration host factor (IHF) in maintenance of plasmid pSC101 in Escherichia coli: mutations in the topA gene allow pSC101 replication in the absence of IHF. Journal of bacteriology 171, 2066–2074.

21. Kasho, K., Fujimitsu, K., Matoba, T., Oshima, T. & Katayama, T. (2014). Timely binding of IHF and Fis to DARS2 regulates ATP–DnaA production and replication initiation. Nucleic acids research 42, 13134–13149.

22. Huang, T., Yuan, H., Fan, L. & Morigen, M. (2020). H-NS, IHF, and DnaA lead to changes in nucleoid organizations, replication initiation, and cell division. Journal of Basic Microbiology 60, 136–148.

23. Dame, R. T., Rashid, F.-Z. M. & Grainger, D. C. (2019). Chromosome organization in bacteria: mechanistic insights into genome structure and function. Nature Reviews Genetics, 1–16.

24. Wei, J., Czapla, L., Grosner, M. A., Swigon, D. & Olson, W. K. (2014). DNA topology confers sequence specificity to nonspecific architectural proteins. Proceedings of the National Academy of Sciences 111, 16742–16747.

25. Ali, B. J., Amit, R., Braslavsky, I., Oppenheim, A. B., Gileadi, O. & Stavans, J. (2001). Compaction of single DNA molecules induced by binding of integration host factor (IHF). Proceedings of the National Academy of Sciences 98, 10658–10663.

26. Azam, T. A., Iwata, A., Nishimura, A., Ueda, S. & Ishihama, A. (1999). Growth phase-dependent variation in protein composition of the Escherichia coli nucleoid. Journal of bacteriology 181, 6361–6370.

27. Makris, J., Nordmann, P. & Reznikoff, W. (1990). Integration host factor plays a role in IS50 and Tn5 transposition. Journal of bacteriology 172, 1368–1373.

28. Wiater, L. & Grindley, N. (1988). Gamma delta transposase and integration host factor bind cooperatively at both ends of gamma delta. The EMBO Journal 7, 1907–1911.

29. Sanyal, S. J., Yang, T.-C. & Catalano, C. E. (2014). Integration host factor assembly at the cohesive end site of the bacteriophage lambda genome: Implications for viral DNA packaging and bacterial gene regulation. Biochemistry 53, 7459–7470.

30. Novotny, L. A., Amer, A. O., Brockson, M. E., Goodman, S. D. & Bakaletz, L. O. (2013). Structural stability of Burkholderia cenocepacia biofilms is reliant on eDNA structure and presence of a bacterial nucleic acid binding protein. PloS one 8, e67629.

31. Gustave, J. E., Jurcisek, J. A., McCoy, K. S., Goodman, S. D. & Bakaletz, L. O. (2013). Targeting bacterial integration host factor to disrupt biofilms associated with cystic fibrosis. Journal of Cystic Fibrosis 12, 384–389.

32. Rao, S. S., Huntley, M. H., Durand, N. C., Stamenova, E. K., Bochkov, I. D., Robinson, J. T., Sanborn, A. L., Machol, I., Omer, A. D. & Lander, E. S. (2014). A 3D map of the human genome at kilobase resolution reveals principles of chromatin looping. Cell 159, 1665–1680.

33. Torbeev, V. Y., Raghuraman, H., Hamelberg, D., Tonelli, M., Westler, W. M., Perozo, E. & Kent, S. B. (2011). Protein conformational dynamics in the mechanism of HIV-1 protease catalysis. Proceedings of the National Academy of Sciences 108, 20982–20987.

34. Sabantsev, A., Levendosky, R. F., Zhuang, X., Bowman, G. D. & Deindl, S. (2019). Direct observation of coordinated DNA movements on the nucleosome during chromatin remodelling. Nature communications 10, 1–12.

35. Porrúa, O., López-Sánchez, A., Platero, A. I., Santero, E., Shingler, V. & Govantes, F. (2013). An A-tract at the AtzR binding site assists DNA binding, inducer-dependent repositioning and transcriptional activation of the PatzDEF promoter. Molecular microbiology 90, 72–87.

36. Corona, T., Bao, Q., Christ, N., Schwartz, T., Li, J. & Dröge, P. (2003). Activation of site-specific DNA integration in human cells by a single chain integration host factor. Nucleic acids research 31, 5140–5148.

37. Kerppola, T. K. & Curran, T. (1991). DNA bending by Fos and Jun: the flexible hinge model. Science 254, 1210–1214.

38. Bera, S. C., Sanyal, K., Senapati, D. & Mishra, P. P. (2016). Conformational Changes Followed by Complete Unzipping of DNA Double Helix by Charge-Tuned Gold Nanoparticles. The Journal of Physical Chemistry B 120, 4213–4220.

39. Jose, D., Weitzel, S. E. & von Hippel, P. H. (2012). Breathing fluctuations in position-specific DNA base pairs are involved in regulating helicase movement into the replication fork. Proceedings of the National Academy of Sciences 109, 14428–14433.

40. Connolly, M., Arra, A., Zvoda, V., Steinbach, P. J., Rice, P. A. & Ansari, A. (2018). Static kinks or flexible hinges: multiple conformations of bent DNA bound to integration host factor revealed by fluorescence lifetime measurements. The Journal of Physical Chemistry B 122, 11519–11534.

41. Velmurugu, Y., Vivas, P., Connolly, M., Kuznetsov, S. V., Rice, P. A. & Ansari, A. (2018). Two-step interrogation then recognition of DNA binding site by Integration Host Factor: an architectural DNA-bending protein. Nucleic acids research 46, 1741–1755.

42. Drozdetski, A. V., Mukhopadhyay, A. & Onufriev, A. V. (2019). Strongly bent double-stranded DNA: reconciling theory and experiment. arXiv preprint 1907.01585.

43. Le, T. T. & Kim, H. D. (2014). Probing the elastic limit of DNA bending. Nucleic acids research 42, 10786–10794.

44. Seong, G. H., Kobatake, E., Miura, K., Nakazawa, A. & Aizawa, M. (2002). Direct atomic force microscopy visualization of integration host factor-induced DNA bending structure of the promoter regulatory region on the Pseudomonas TOL plasmid. Biochemical and biophysical research communications 291, 361–366.

45. Beckwitt, E. C., Kong, M. & Van Houten, B. (2018). Seminars in cell & developmental biology.

46. Thomson, N. H., Santos, S., Mitchenall, L. A., Stuchinskaya, T., Taylor, J. A. & Maxwell, A. (2014). DNA G-segment bending is not the sole determinant of topology simplification by type II DNA topoisomerases. Scientific reports 4, 1–9.

47. Koonin, E. V. (2014). Carl Woese’s vision of cellular evolution and the domains of life. rNa Biology 11, 197–204.

48. Theobald, D. L. (2010). A formal test of the theory of universal common ancestry. Nature 465, 219–222.

49. Doolittle, W. F. (2000). Uprooting the tree of life. Scientific American 282, 90–95.

50. Glansdorff, N., Xu, Y. & Labedan, B. (2008). The last universal common ancestor: emergence, constitution and genetic legacy of an elusive forerunner. Biology direct 3, 29.

51. Bakshi, S., Siryaporn, A., Goulian, M. & Weisshaar, J. C. (2012). Superresolution imaging of ribosomes and RNA polymerase in live Escherichia coli cells. Molecular microbiology 85, 21–38.

52. Manning, G. S. (2006). The persistence length of DNA is reached from the persistence length of its null isomer through an internal electrostatic stretching force. Biophysical journal 91, 3607–3616.

53. Koo, H.-S. & Crothers, D. M. (1988). Calibration of DNA curvature and a unified description of sequence-directed bending. Proceedings of the National Academy of Sciences 85, 1763–1767.

54. Bolshoy, A., McNamara, P., Harrington, R. & Trifonov, E. (1991). Curved DNA without AA: experimental estimation of all 16 DNA wedge angles. Proceedings of the National Academy of Sciences 88, 2312–2316.

55. Connolly, M., Zvoda, V. & Ansari, A. (2018). Equilibrium Conformational Distributions of Bent DNA in Complex with IHF Mapped with Fluorescence Lifetime Measurements. Biophysical Journal 114, 29a–30a.

56. Nagaich, A. K., Zhurkin, V. B., Durell, S. R., Jernigan, R. L., Appella, E. & Harrington, R. E. (1999). p53-induced DNA bending and twisting: p53 tetramer binds on the outer side of a DNA loop and increases DNA twisting. Proceedings of the National Academy of Sciences 96, 1875–1880.

57. Fei, J., Torigoe, S. E., Brown, C. R., Khuong, M. T., Kassavetis, G. A., Boeger, H. & Kadonaga, J. T. (2015). The prenucleosome, a stable conformational isomer of the nucleosome. Genes & development 29, 2563–2575.

58. Swinger, K. K. & Rice, P. A. (2007). Structure-based analysis of HU–DNA binding. Journal of molecular biology 365, 1005–1016.

59. Dame, R. T., Noom, M. C. & Wuite, G. J. (2006). Bacterial chromatin organization by H-NS protein unravelled using dual DNA manipulation. Nature 444, 387–390.

60. Dame, R. T., Wyman, C. & Goosen, N. (2000). H-NS mediated compaction of DNA visualised by atomic force microscopy. Nucleic acids research 28, 3504–3510.

61. Zhang, N., Kuznetsov, S. G., Sharan, S. K., Li, K., Rao, P. H. & Pati, D. (2008). A handcuff model for the cohesin complex. The Journal of cell biology 183, 1019–1031.

62. Das, A., Chatterjee, S. & Raghuraman, H. (2020). Structural dynamics of the paddle motif loop in the activated conformation of KvAP voltage sensor. Biophysical journal 118, 873–884.

63. Maria-Solano, M. A., Serrano-Hervás, E., Romero-Rivera, A., Iglesias-Fernández, J. & Osuna, S. (2018). Role of conformational dynamics in the evolution of novel enzyme function. Chemical Communications 54, 6622–6634.

64. Adhya, S. (1989). Multipartite genetic control elements: communication by DNA loop. Annual review of genetics 23, 227–250.

65. Malik, M., Bensaid, A., Rouviere-Yaniv, J. & Drlica, K. (1996). Histone-like protein HU and bacterial DNA topology: suppression of an HU deficiency by gyrase mutations. Journal of molecular biology 256, 66–76.

66. Rouvière-Yaniv, J., Yaniv, M. & Germond, J.-E. (1979). E. coli DNA binding protein HU forms nucleosome-like structure with circular double-stranded DNA. Cell 17, 265–274.

67. Zhang, N., Darbari, V. C., Glyde, R., Zhang, X. & Buck, M. (2016). The bacterial enhancer-dependent RNA polymerase. Biochemical Journal 473, 3741–3753.

68. Cruz-Becerra, G. & Kadonaga, J. T. (2020). Enhancement of homology-directed repair with chromatin donor templates in cells. Elife 9, e55780.

69. Alexiadis, V. & Kadonaga, J. T. (2002). Strand pairing by Rad54 and Rad51 is enhanced by chromatin. Genes & development 16, 2767–2771.

70. Bao, Q., Christ, N. & Dröge, P. (2004). Single-chain integration host factors as probes for high-precision nucleoprotein complex formation. Gene 343, 99–106.

71. Aitken, C. E., Marshall, R. A. & Puglisi, J. D. (2008). An oxygen scavenging system for improvement of dye stability in single-molecule fluorescence experiments. Biophysical journal 94, 1826–1835.

72. Paul, T., Bera, S. C., Agnihotri, N. & Mishra, P. P. (2016). Single-molecule FRET studies of the hybridization mechanism during noncovalent adsorption and desorption of DNA on graphene oxide. The Journal of Physical Chemistry B 120, 11628–11636.

73. McKinney, S. A., Joo, C. & Ha, T. (2006). Analysis of single-molecule FRET trajectories using hidden Markov modeling. Biophysical journal 91, 1941–1951.

74. Lee, T.-H. (2009). Extracting kinetics information from single-molecule fluorescence resonance energy transfer data using hidden Markov models. The Journal of Physical Chemistry B 113, 11535–11542.

75. Šulc, P., Romano, F., Ouldridge, T. E., Rovigatti, L., Doye, J. P. & Louis, A. A. (2012). Sequence-dependent thermodynamics of a coarse-grained DNA model. The Journal of chemical physics 137, 135101.

76. Blender, O. (2018). Blender—A 3D modelling and rendering package. Retrieved.

