## Supplementary material for "Vital Insights into Prokaryotic Genome Compaction by Nucleoid-Associated Protein (NAP) and Illustration of DNA Flexure Angles at Single Molecule Resolution": IHF_Supplementary

### * Corresponding author

### Supporting Information

### DNA Constructs:

### Labeling Scheme in DNA-Protein Complex:

Four DNA constructs having similar sequence but shifts in position of the labeled donor and acceptor has been used in this study as shown in the figure. The sequence consists of A-tract (6A repeats) region and H’-binding sequence (TATCAA) for IHF.

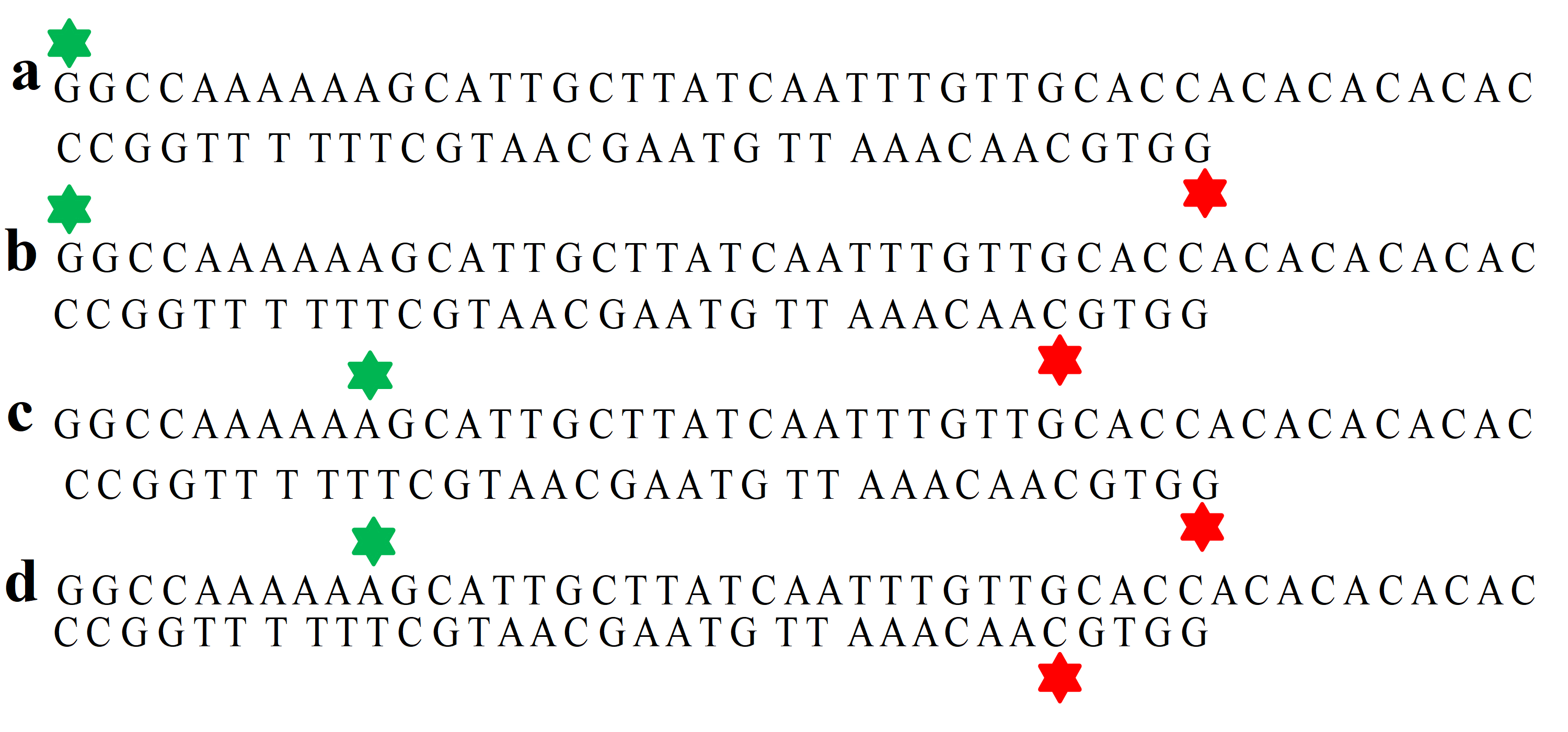

**Figure S1.** The points of labeling with the donor and acceptor in the four DNA constructs have been represented. **(a)**Represents Construct 1 where donor and acceptor are at the end. **(b)** The acceptor is shifted five nucleotides from the end and donor is at the terminal in Construct 2. **(c)** Donor is shifted 10 nucleotides from the terminal and acceptor is at the end of sequence in Construct 3. **(d)** In Construct 4 both the donor and acceptor are shifted 10 and 5 nucleotides from the end

**TableS1.**List of sequences for all the oligonucleotides used in this study. Modifications are mentioned along with the sequences, with specific color coding. All the oligoes are HPLC purified and purchased from IDT.

| Sequence Name | Sequence & Modification (5’-3’) |
| --- | --- |
| Sequence 1 | **(Cy3)**GGCCAAAAAAGCATTGCTTATCAATTTGTTGCACCACACACACAC |
| Sequence 2 | GGCCAAAAAA**(Cy3)**GCATTGCTTATCAATTTGTTGCACCACACACACAC |
| Sequence 3 | **(Cy5)**GGTGCAACAAATTGATAAGCAATGCTTTTTTGGCC |
| Sequence 4 | GGTGC**(Cy5)**AACAAATTGATAAGCAATGCTTTTTTGGCC |
| Sequence 5 | **BIOTIN**-AAAAATGTGTGTGTG |

### Representative smFRET traces of Naked DNA:

Time traces of four constructs of naked DNA (only DNA without protein). All the molecules display low FRET state, 0.1 in case of Construct1, 2 and 3. The average FRET Efficiency of Construct 4 comes out as 0.2.

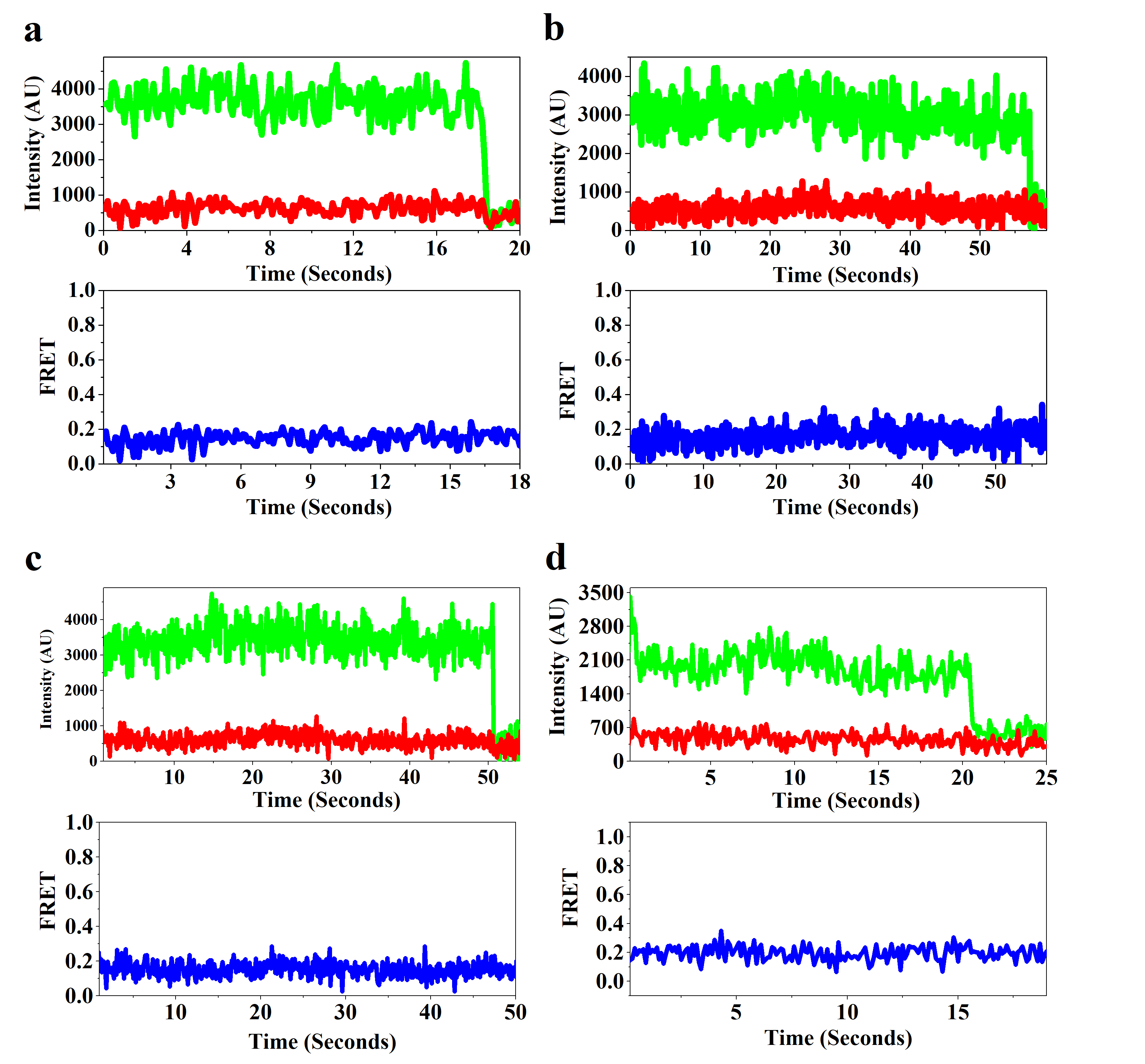

**Figure S2.** Time traces and FRET Efficiency of representative single molecules corresponding to the four constructs of DNA **(a)** to **(d)** stands for Construct1 to 4 sequentially

**Parameters from Crystal Structure**

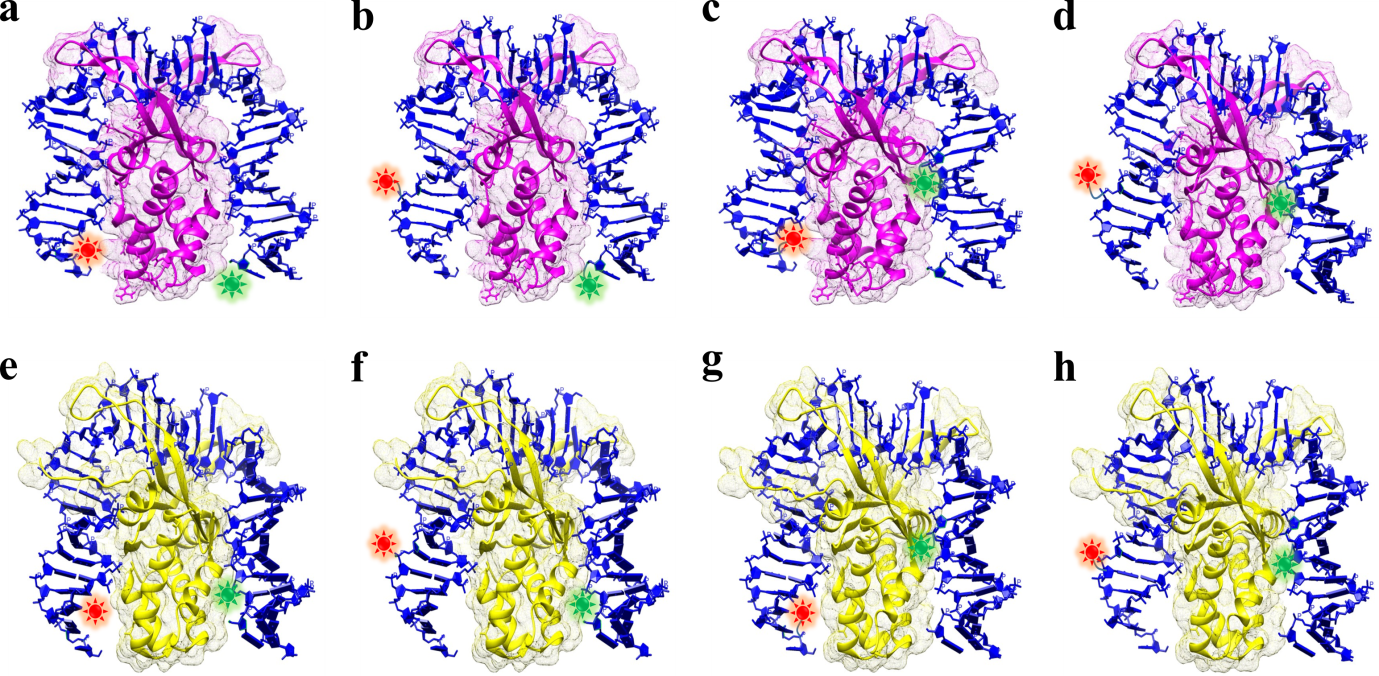

**Figure S3** Representative models of the crystal structure, with green and red fluorophore depicting donor and acceptor positions respectively in the DNA. The protein wtIHF(PDBID: 1IHF)is shown in purple colour and scIHF (PDBID: 2IIE) is represented in yellow colour. **(a)**to**(d)** and **(e)** to **(h)** corresponds to Construct 1 to 4 in order with respect to wtIHF and scIHF. The models has been made using Chimera 1.13.1rc.

The expected distance between the dyes in all the four DNA constructs if the DNA attains the conformation of the crystal structure upon binding with both scIHF and wtIHF, has been computed. Also, the corresponding FRET Efficiencies have been listed.

**Table S2: Distances corresponding to Cy3 & Cy5 labelling for scIHF&wtIHF (corresponding FRET efficiencies are given in the box bracket)**

|  | **Atom Specifier Cy3** | **Atom Specifier Cy5** | **scIHF (2IIE.PDB)** | **wtIHF (1IHF.PDB)** |
| --- | --- | --- | --- | --- |
| **Combo1** | 15.D O5’ | -49.C P | 40.124 A [0.85] | 39.144 A [0.87] |
| **Combo2** | 15.D O5’ | -45.C P | 45.764 A [0.75] | 45.981 A [0.75] |
| **Combo3** | 24.D P | -49.C P | 38.826 A [0.89] | 39.388 A [0.87] |
| **Combo4** | 24.D P | -45.C P | 43.364 A [0.79] | 43.444 A [0.79] |

### Geometry for Flexure Angle Determination:

**A**

**B (0,0)**

**D_pro_**

**C (x,0)**

**X**

**θ**

**B’**

**C’**

**α+α_0_**

**β+ β_0_**

**r**

**R**

**Z**

**Figure S4.**Quadrilateral, ABCD depicting the outline of bent DNA. B has been taken as the origin, and the coordinates of C are (x,0). Length of AB is **‘r’**and length of CD is ‘**R**’. As per the coordinates, length of BC is **x**. X is the mid point of BC. AD_pro_ length is taken as Z.

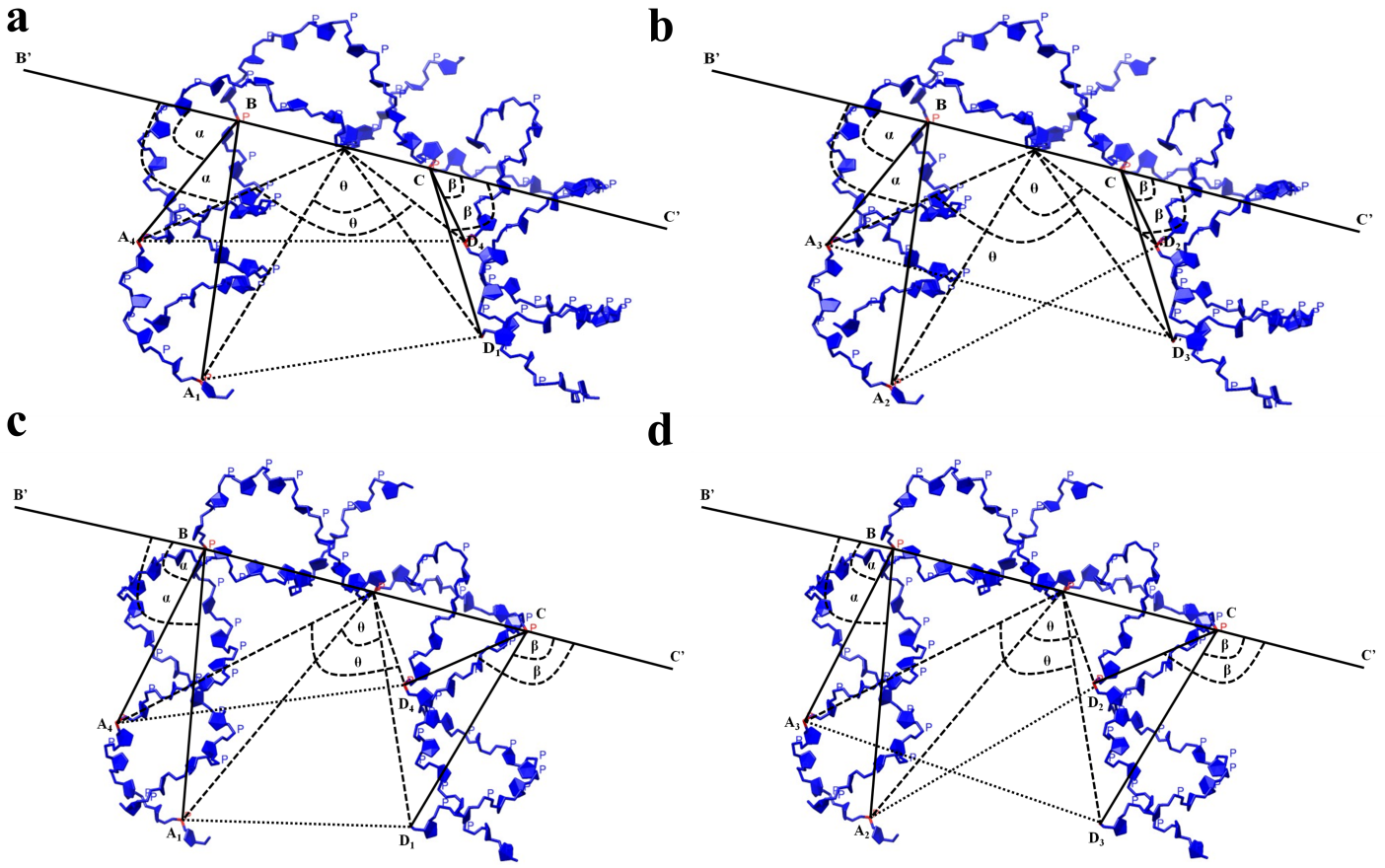

**Figure S5** Outline of the DNA construct, established in quadrilateral model. **(a)**and**(b)**, depicts outline of DNA Constructs 1, 4 and 2, 3 respectively bound with wtIHF**(c)** and **(d)**follows the same order of DNA constructs bound with scIHF. The models has been made using Chimera 1.13.1rc.

*A = (rcos(Π-(α+α_0_)), rsin(Π-(α+α_0_)))*

*or, A = (-rcos(α+α_0_), rsin(α+α_0_)*

*D_pro_= (x-Rcos(Π-(β+β_0_)), Rsin(Π-(β+β_0_)))*

*or, D_pro_ = (x+Rcos(β+β_0_), Rsin(β+β_0_))*

***Z^2^ = [x + Rcos(β+β_0_) + rcos(α+α_0_)]^2^ + [Rsin(β+β_0_)-rsin(α+α_0_)]^2^……………………(1)***

***XA^2^ = [l+ rcos(α+α_0_)]^2^ + [rsin(α+α_0_)]^2^ = a^2^……………………………(2)***

***XD_pro_^2^ = [x+Rcos(β +β_0_) - l]^2^ + [Rsin(β+β_0_)]^2^ = b^2^…………………….(3)***

*cosθ = (a^2^ + b^2^ – Z^2^)/2a*b*

***θ = cos^-1^[(a^2^+b^2^– Z^2^)/2a*b]……………………………...(4)***

Since, binding of both the proteins introduce two kinks in the DNA (scIHF and wtIHF), to mimic a regular structure the outline of DNA has been exemplified as a quadrilateral. The vertices, B and C of the quadrilateral are the points in the DNA where protein induces the kink. X is taken as the midpoint of the arm BC. The vertices A and D are the ends of DNA whose positions are changed according to the point of fluorophore labelling. The lengths AB, BC and CD are found using the vector coordinates from the crystal structure. The whole picture comes out in three dimension, but inspite of calculating the solid flexure angle, which can increase the number of variables, hence, more set of assumptions and finally a very approximated value of the flexure angle, we have chosen to find the two dimensional angle by acquiring a projection of the arm CD on the plane containing AB and BC, which is CD_pro_.

The vector coordinates of A, B, C and D has been determined from the crystal structure of the DNA bound with wtIHF and scIHF (PDBID: 1IHF and 2IIE). In this study, in order to bring more accuracy for finding DNA bending parameters, angle AXD_pro_ termed as flexure angle and represented as **‘θ’** is determined. The coordinates of D_pro_ has been determined by taking the projection of arm CD on the plane containing A, B and C so that, we can get a 2 dimensional projection of the whole system. Upon referring to the reports from the crystal structure it was found that, the angle between the arm CD and the plane having AB and BC, is very small. So, considering the whole picture in two dimensions does not deviate much from the original three dimensional arrangement.α and β are the deviation from external angles, (**α_0_, β_0_**) of the quadrilateral between the sides BC, BA and CB, CD_pro_ respectively whose initial values are 0 degree, calculated to the corresponding crystal structures. The length AD_pro_ (Z) is determined using the FRET Efficiency value and feeding it into the equation, E_FRET_=1/ (1+(R/R_0_)^6^) where R_0_ is taken as 5.4 nm. Since, our experimental results shows a dilated conformation of bent state of the DNA than that of the crystal structure, the range of α and β has been varied as a set of angles from [0,**-α_0_**] and [0,**-β_0_**]. Now, the keeping the Z-value fixed (as obtained from smFRET experiments), a set of α and β values has been generated satisfying the experimental Z-value by feeding them in equation (1). Using, the values of determined α and β, *a* and *b* values are calculated through equation (2) and (3) respectively. And finally, the set of θ-values are determined using equation (4). Even though we got a family of solution for θ by applying the above mentioned mechanism for a particular Z-value, the distribution did not vary much ranging between 1 to 6 degrees. The programme was written and executed on Mathematica.

### Table S3: α_0_ and β_0_ for different constructs (Value from the quadrilateral extracted from crystal structure):

| **Constructs** | **α_0_ (degree)** | **β_0_ (degree)** |
| --- | --- | --- |
| C1wtIHF | 97.48 | 68.67 |
| C2wtIHF | 75.14 | 71.84 |
| C3wtIHF | 97.48 | 59.75 |
| C4wtIHF | 75.14 | 62.47 |
| C1scIHF | 98.48 | 69.29 |
| C2scIHF | 76.06 | 73.13 |
| C3scIHF | 98.48 | 59.63 |
| C4scIHF | 76.06 | 63.19 |

### 3-Dimensional plots:

As mentioned above, in order to find the flexure angle, the first step is to generate the set of angular values of α and β, to get the value of θ (flexure angle). The values of α and β so found and the corresponding flexure angle values have been plotted in a 3-dimensional plot where x, y and z-axis represents corresponding values of α, β and θ respectively.

### wtIHF

**a**
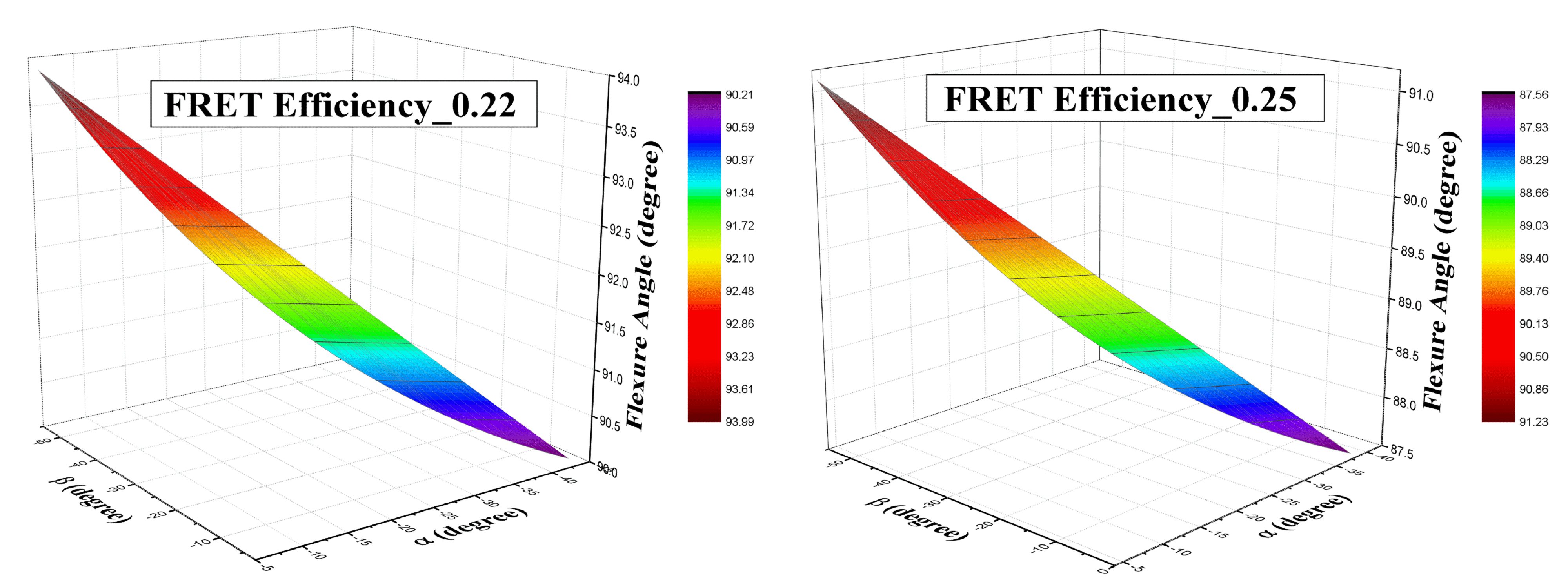

**b**
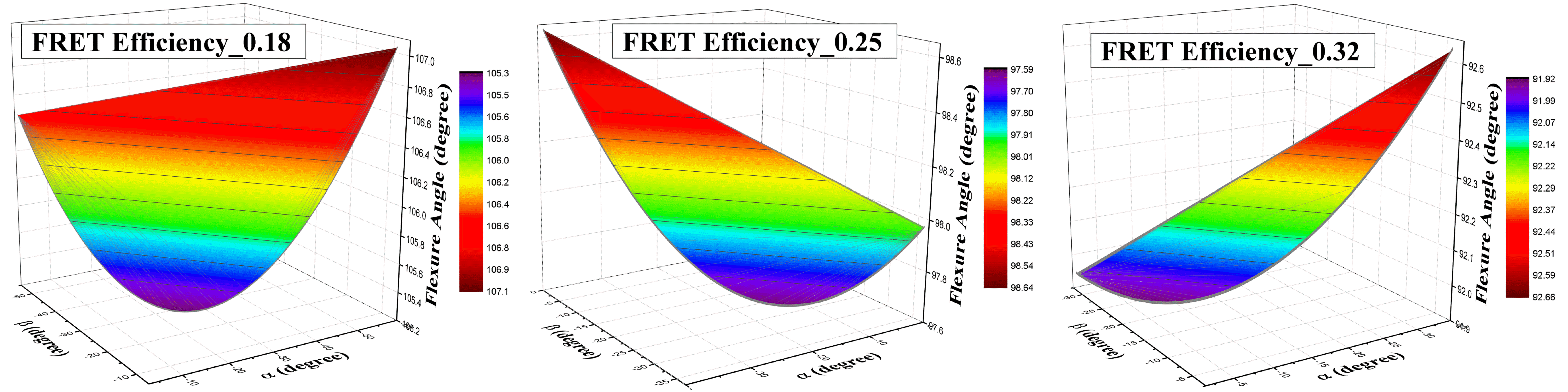

**c**
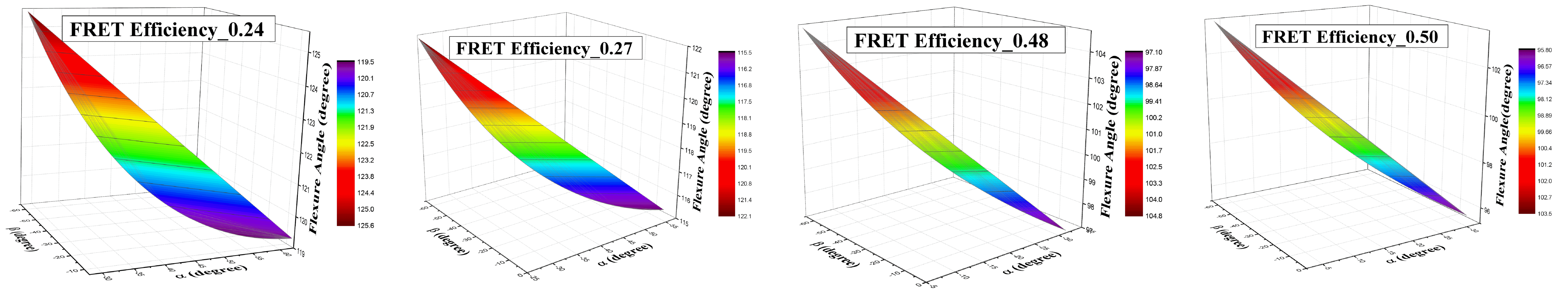

**d**
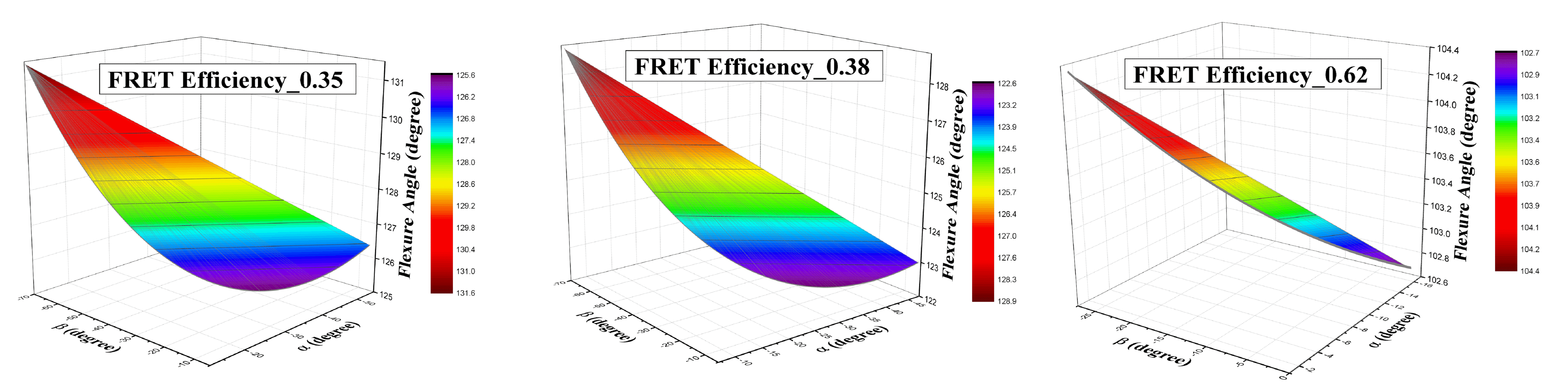

**Figure S6.** Graphs in section **(a), (b), (c)** and **(d)** are the plots of angular values when constructs 1 to 4 of DNA is subjected to bind with wtIHF. The FRET Efficiency values are mentioned in the boxes.

**scIHF**

**a**
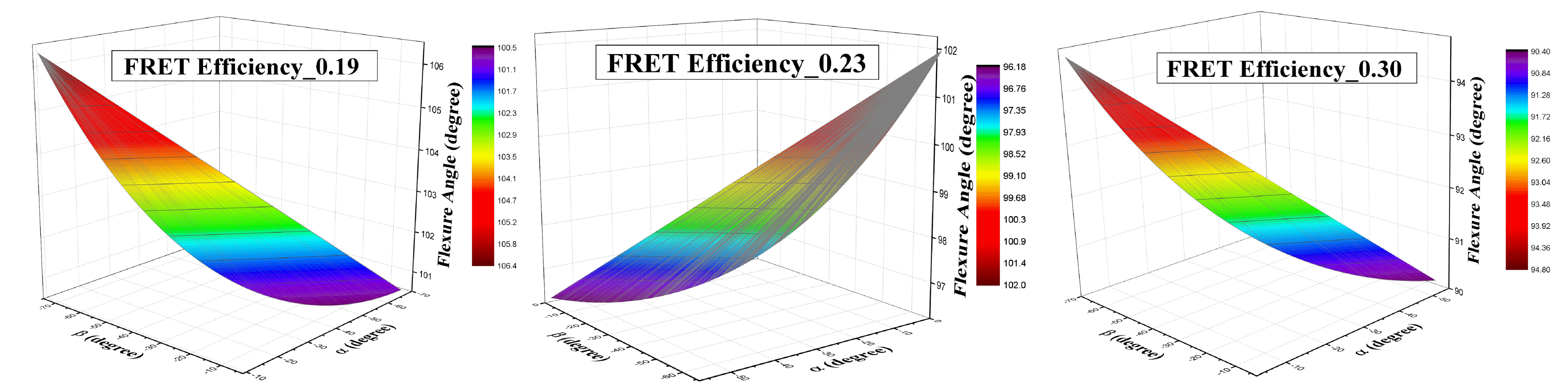

**b**
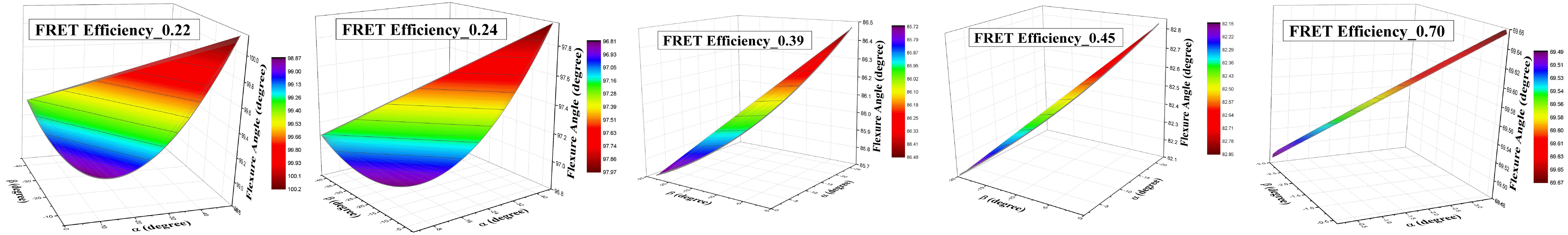

**c**
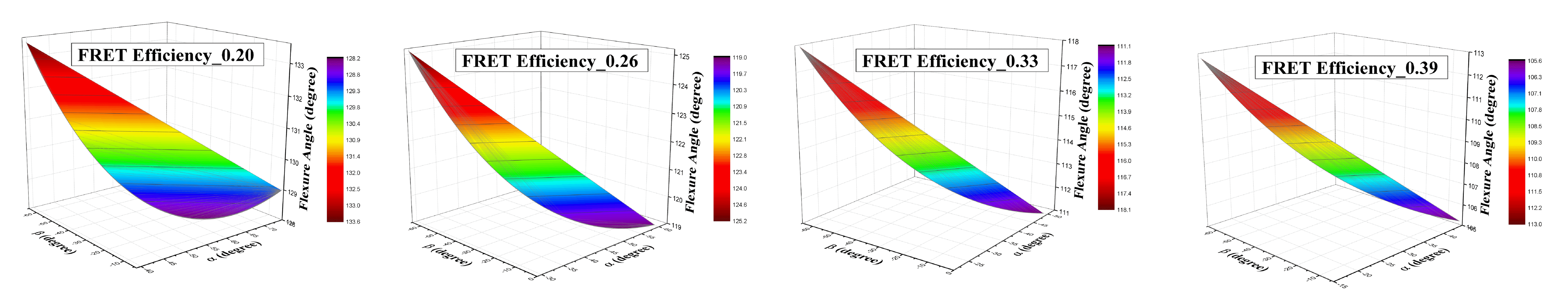

**d**
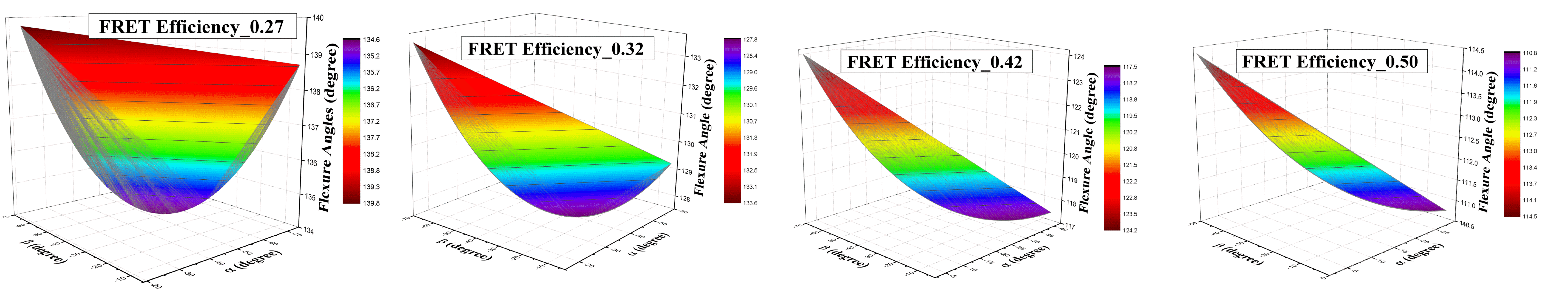

**Figure S7.** At eachFRET Efficiency for each construct upon scIHF binding, the correlated values of α, β and θ (flexure angle) are plotted in a 3-dimensional plot. **(a)**to**(d)** stands sequentially for Construct 1 to 4.

### Histograms:

The set of flexure angle values so found plotted in the form of histogram, with number of occurrences in the y-axis and angular values in the x-axis. This gives us the idea of the distribution probability at each angular value in the range of angles so found.

**wtIHF**

**a**
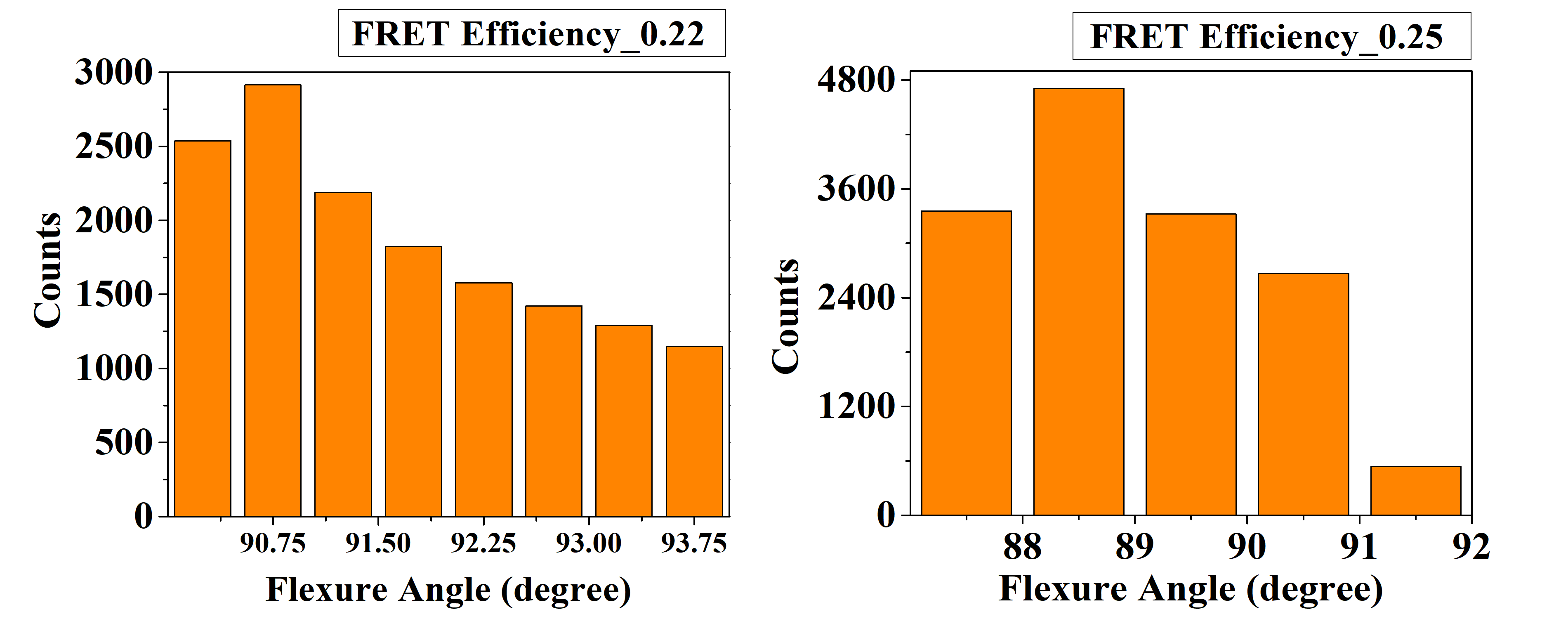

**b**
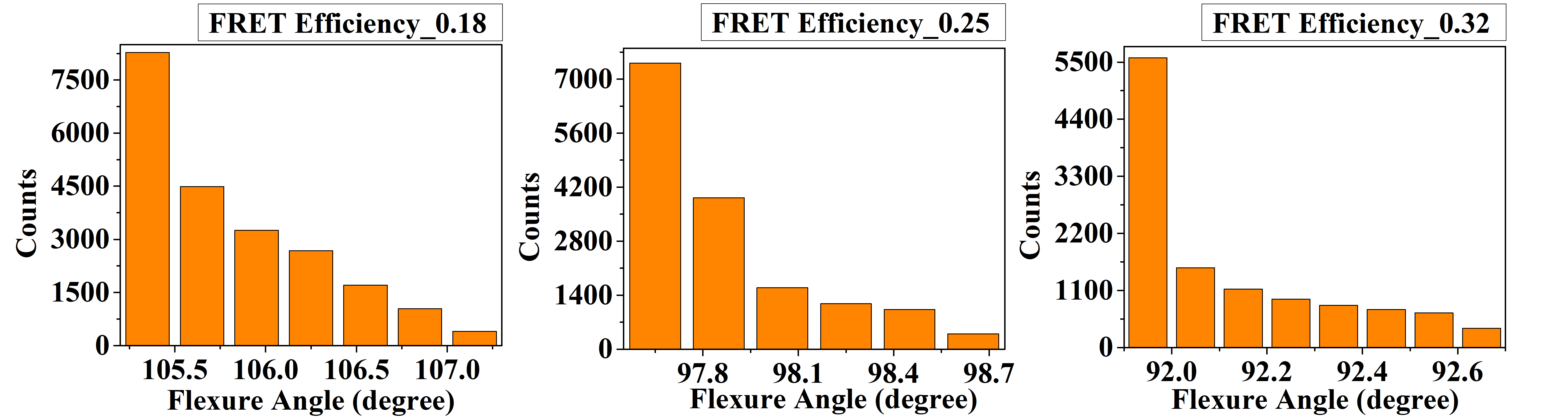
 **c**
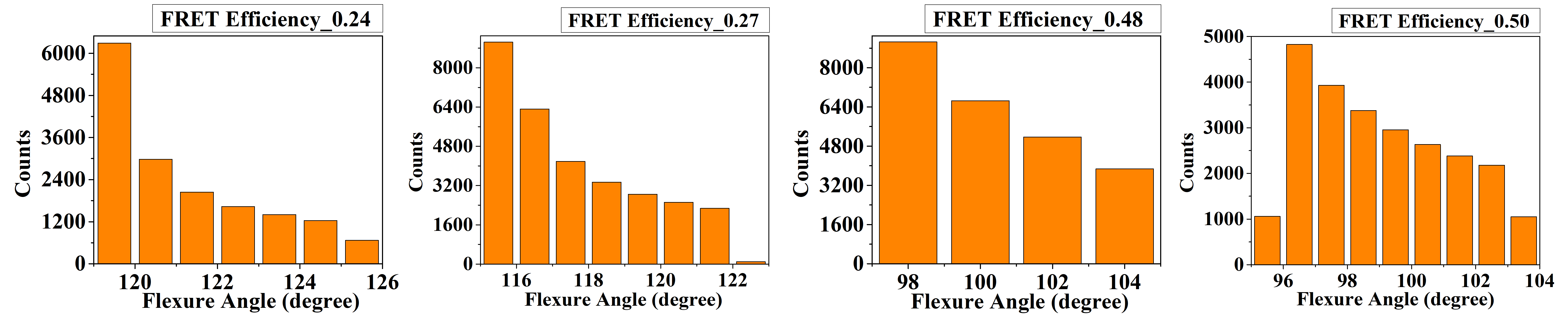

**d**
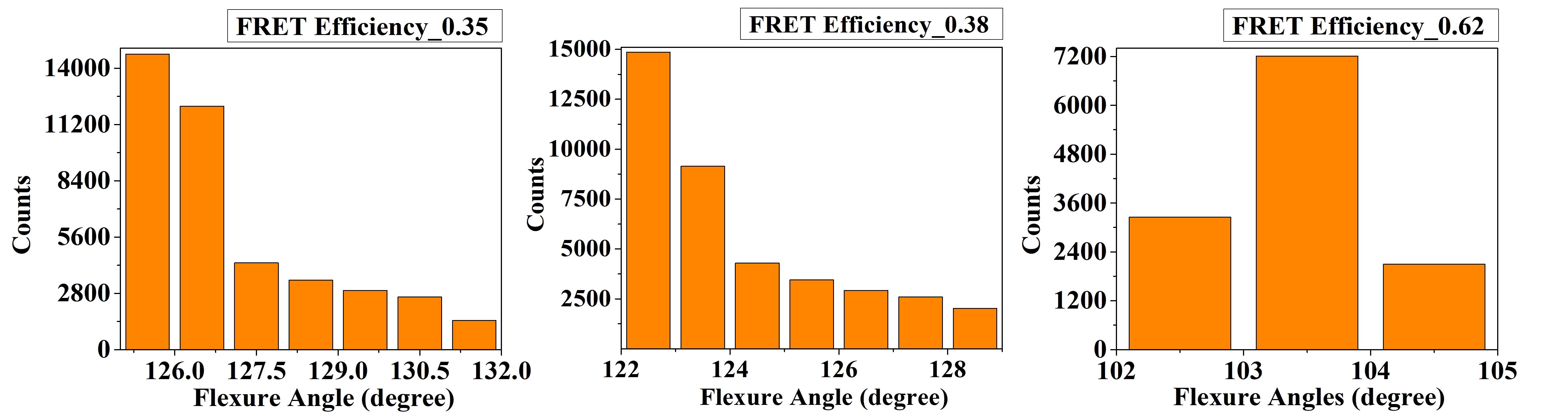

**Figure S8. (a)** to**(d)** sequentially illustrates flexure angle values for constructs 1 to 4 when they are bound with wtIHF. The FRET Efficiency values are mentioned at the top of the plots.

**scIHF**

**a**
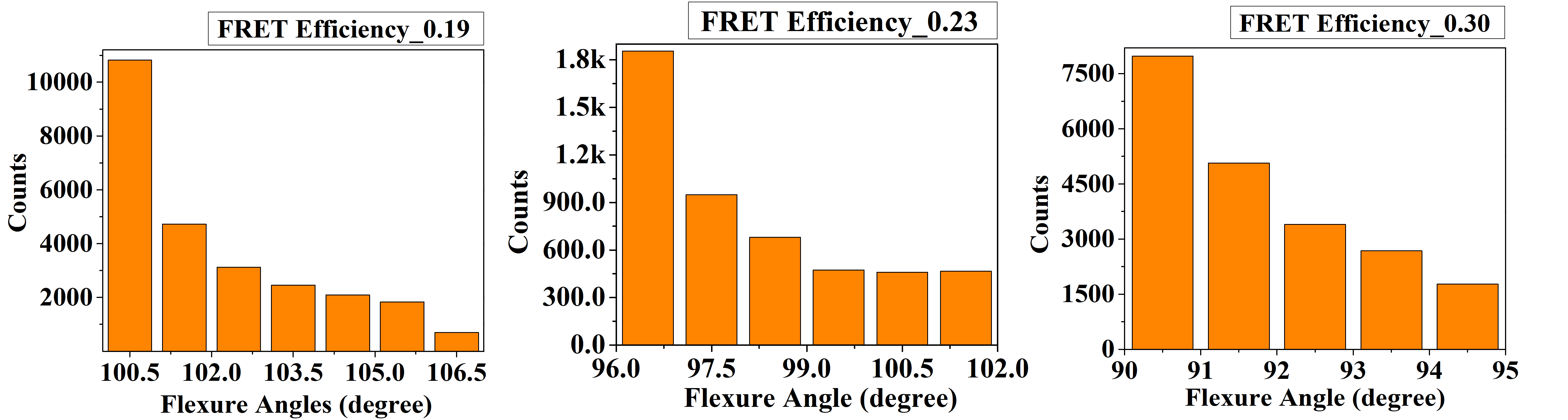

**b**
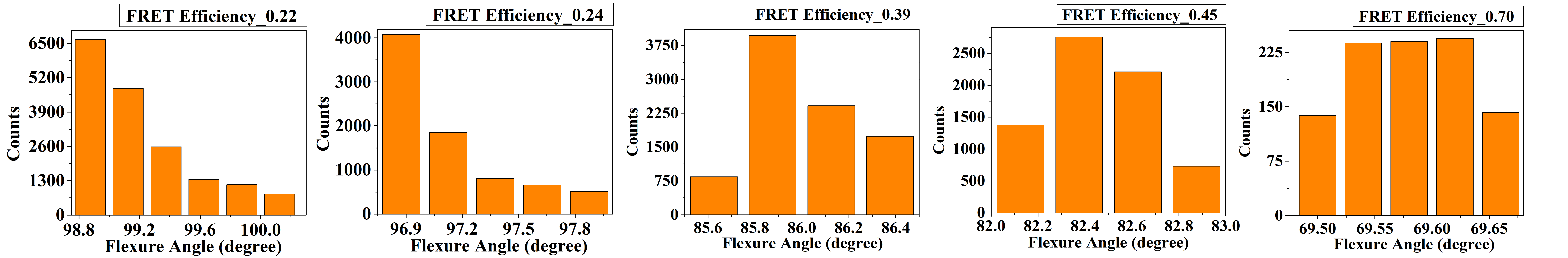
 **c**
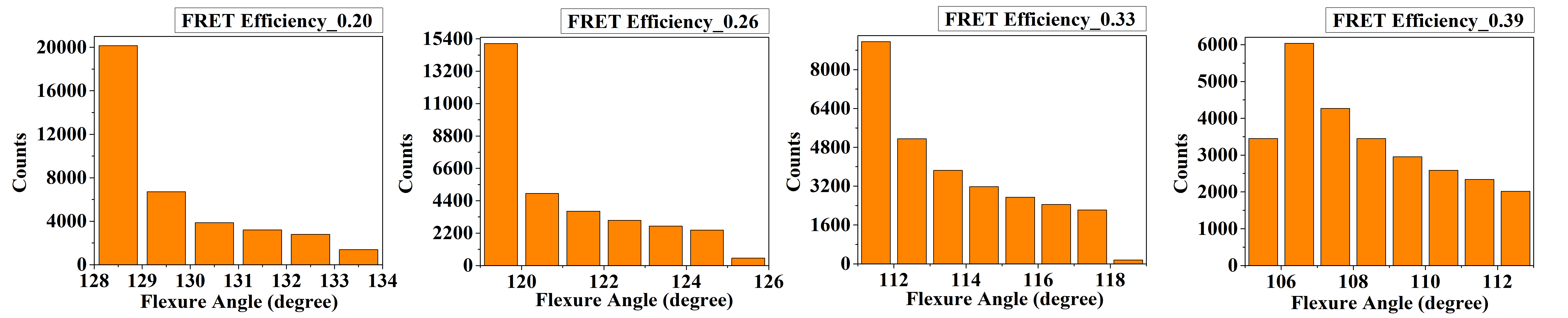

**d**
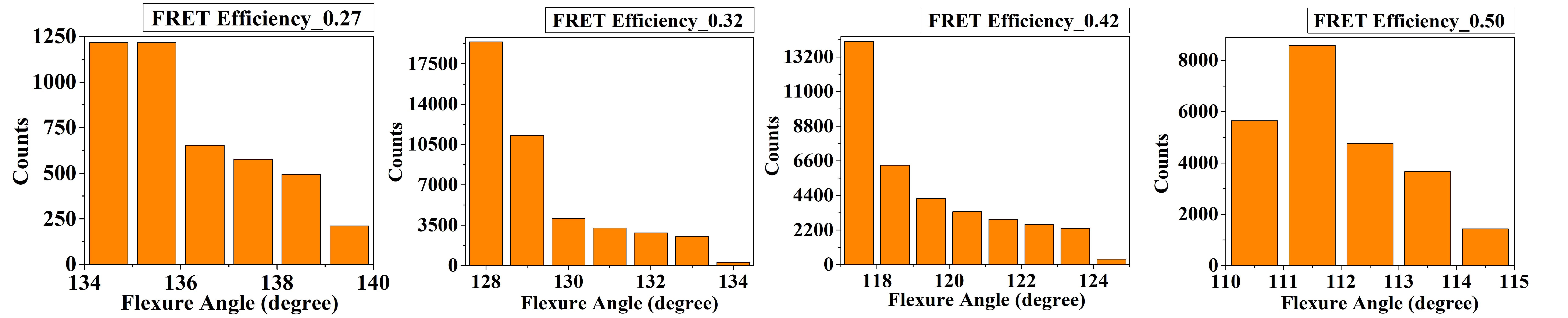

**Figure S9.** Representation of the flexure angle histogram, in figure subset **(a)** to **(d)** when DNA constructs 1 to 4 are allowed to bind with scIHF.

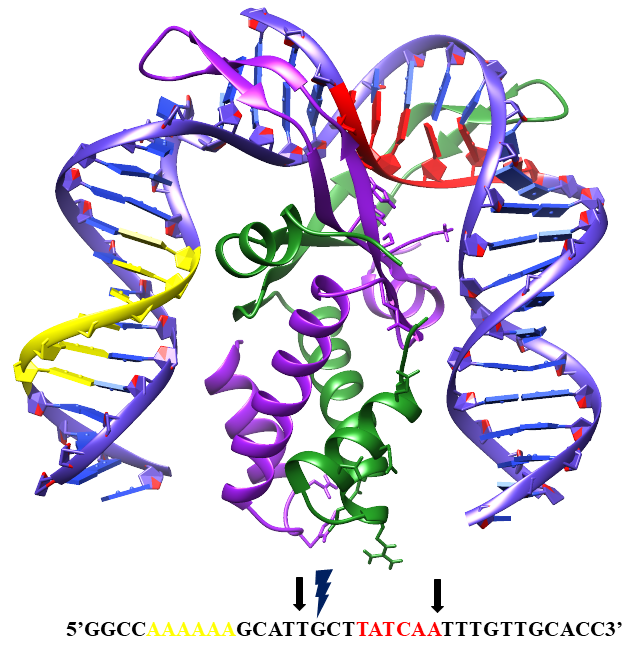

**Figure S10.** Representation the IHF-DNA complex (PBD ID: 1IHF). The two subunits of the IHF, α and β has been highlighted in Green and Purple respectively. The A-tract region and the H’ sequence has been highlighted in Yellow and Red respectively. The DNA sequence has been given at the bottom, the two Black arrows indicate the positions of the kink by IHF and the Thunderbolt indicates the nick. The model has been made using Chimera 1.13.1rc.

**Table S4:** Mean flexure angle values and Standard Deviation (S.D) corresponding to the FRET Efficiency in degree upon binding of DNA with wtIHF, **(a)** to **(d)** stand for Construct 1 to 4 in order.

**(a)**

| **FRET Efficiency** | **Mean Flexure Angle (degree)** | **Std. Dev.** |
| --- | --- | --- |
| 0.22 | 91.7 | 1.1 |
| 0.25 | 89.0 | 1.1 |

**(b)**

| **FRET Efficiency** | **Mean Flexure Angle (degree)** | **Std. Dev.** |
| --- | --- | --- |
| 0.18 | 105.8 | 0.5 |
| 0.25 | 97.9 | 0.3 |
| 0.32 | 92.1 | 0.2 |

**(c)**

| **FRET Efficiency** | **Mean Flexure Angle (degree)** | **Std. Dev.** |
| --- | --- | --- |
| 0.24 | 121.2 | 1.8 |
| 0.27 | 117.6 | 1.9 |
| 0.48 | 100.3 | 2.2 |
| 0.50 | 99.0 | 2.2 |

**(d)**

| **FRET Efficiency** | **Mean Flexure Angle (degree)** | **Std. Dev.** |
| --- | --- | --- |
| 0.35 | 127.1 | 1.7 |
| 0.38 | 124.2 | 1.8 |
| 0.62 | 103.4 | 0.5 |

**Table S5:** Mean values of flexure angle and standard deviation with respect to the FRET Efficiency when the DNA is allowed to bind with scIHF, **(a)** to **(d)** are in the order from Construct 1 to 4.

**(a)**

| **FRET Efficiency** | **Mean Flexure Angle (degree)** | **Std. Dev.** |
| --- | --- | --- |
| 0.19 | 102.1 | 1.7 |
| 0.23 | 98.1 | 1.8 |
| 0.30 | 91.8 | 1.3 |

**(b)**

| **FRET Efficiency** | **Mean Flexure Angle (degree)** | **Std. Dev.** |
| --- | --- | --- |
| 0.22 | 99.2 | 0.3 |
| 0.24 | 97.1 | 0.3 |
| 0.39 | 86.0 | 0.2 |
| 0.45 | 82.5 | 0.2 |
| 0.70 | 69.6 | 0.1 |

**(c)**

| **FRET Efficiency** | **Mean Flexure Angle (degree)** | **Std. Dev.** |
| --- | --- | --- |
| 0.20 | 129.6 | 1.5 |
| 0.26 | 120.9 | 1.8 |
| 0.33 | 113.6 | 2.0 |
| 0.39 | 108.4 | 2.1 |

**(d)**

| **FRET Efficiency** | **Mean Flexure Angle (degree)** | **Std. Dev.** |
| --- | --- | --- |
| 0.27 | 136.2 | 1.5 |
| 0.32 | 129.3 | 1.6 |
| 0.42 | 119.4 | 1.9 |
| 0.50 | 112.0 | 1.1 |
